# Individual differences in frontal midline theta activity during visuomotor adaptation are related to execution noise

**DOI:** 10.1101/2020.07.12.188581

**Authors:** Zeb D. Jonker, Rick van der Vliet, Guido Maquelin, Joris van der Cruijsen, Gerard M. Ribbers, Ruud W. Selles, Opher Donchin, Maarten A. Frens

## Abstract

Frontal midline EEG activity has been found to correlate with error magnitude during motor adaptation. We replicated a previous visuomotor adaptation experiment with very small perturbations, likely to invoke implicit adaptation, in a new group of 60 participants and combined it with EEG recordings. We used this data to explore 1) whether frontal midline activity will be evoked in the absence of awareness of the perturbation; 2) whether frontal midline activity is related to implicit adaptation; 3) whether individual differences in frontal midline activity are related to individual differences in motor learning. The results showed that frontal midline theta activity (FMΘ) is also present during small perturbations, does not drive between-trial error correction, and that the sensitivity of FMΘ to error magnitude was smaller for participants with greater execution noise. This relation between FMΘ-error-sensitivity and execution noise could be fully explained by looking at the relationship between FMΘ and error probability. This implies that frontal midline theta activity represents a surprise-like saliency signal, potentially driving awareness and cognitive control in situations with more salient errors.

## INTRODUCTION

In motor learning, movement errors are used to update motor commands for subsequent actions. We call this process motor adaptation. Motor adaptation can occur within a trial during continuous movements, or between trials during discrete ballistic movements such as swinging a golf club or throwing darts. In motor adaptation tasks, perturbations of visual or proprioceptive feedback drive adjustments of the motor command to achieve the intended goal of the movement. These adjustments may reflect unconscious implicit adaptation, conscious explicit strategy, or both (Taylor et al., 2014). Typically, smaller gradual perturbations result in implicit adaptation whereas larger sudden perturbations have been found to induce explicit strategy (Malfait and Ostry, 2004; Werner et al., 2019).

Linear dynamical models of error-based learning that include an adaptation rate, a retention rate, planning noise (state noise), and execution noise (output noise) have been shown to qualitatively resemble between-trial implicit adaptation of reaching movements to a gradual perturbation signal with incremental perturbation steps (Cheng and Sabes, 2007). In this model class, planning noise represents stochastic noise in central movement planning, whereas execution noise represents stochastic noise in peripheral movement execution that is not included in the motor command (van Beers, 2009). Van der Vliet et al. (2018) showed that such a model can also explain differences in motor learning between individuals. In that study we presented a novel Bayesian fitting procedure to estimate the model parameters for each individual participant. In agreement with optimal Kalman filter theory (Kalman, 1960), individuals with relatively more planning noise and less execution noise showed a higher adaptation rate than individuals with relatively less planning noise and more execution noise. However, an important question regarding these mathematical models is the extent to which they reflect the reality of the underlying neurophysiological processes (Tan et al., 2016).

Independently from the mathematical modelling approach, electroencephalography (EEG) has been used increasingly to explore neurophysiological correlates of learning. Most studies have focused on frontal midline activity, which has been found to correlate with error magnitude in motor adaptation tasks (Krigolson et al., 2008; Anguera et al., 2009; Vocat et al., 2011; Torrecillos et al., 2014; Maclean et al., 2015; Arrighi et al., 2016; Reuter et al., 2018). However, a recent study with very small gradual perturbations did not find a relation between error magnitude and frontal midline activity (Palidis et al., 2019). Furthermore, to our knowledge, no prior studies have tested whether frontal midline activity was driving adaptation in the consecutive trial.

The current study therefore explores 1) whether frontal midline activity will be evoked in the absence of awareness of the perturbation; 2) whether frontal midline activity is related to implicit adaptation; 3) whether individual differences in frontal midline activity are related to individual differences in motor learning. To achieve these objectives, we replicated our previous visuomotor adaptation experiment (van der Vliet et al., 2018) in a new group of 60 participants and combined it with EEG recordings.

## METHODS

### Participants

We recruited 60 right-handed (Oldfield, 1971) participants (19 men and 41 women; mean age = 25.6 years, range = 18-61) without any medical condition that might interfere with motor performance. Participants received a small financial token for traveling and time compensation. The study was performed in accordance with the Declaration of Helsinki and was approved by the medical ethics committee of the Erasmus MC University Medical Centre.

### Experimental procedure

The experimental procedure was adapted from van der Vliet et al. (2018). Participants were seated in front of a horizontal projection screen with a robotic handle underneath. This handle was situated at elbow height and could be moved in the horizontal plane. The position of the handle was projected on top of the screen as a cursor (green circle 5mm diameter). Furthermore, the screen displayed an origin (white circle 10mm diameter) and a target (red circle 10mm diameter) at fixed positions. The origin was located approximately 40cm straight in front of the participant and the target was projected exactly 10cm behind the origin, approximately 50cm in front of the participant. Participants were instructed to hold the handle in their dominant right hand and move the cursor from the origin through the target in one smooth ballistic reaching movement. To prevent direct feedback of hand position underneath the screen, participants were wearing an apron that was secured to the top of the screen.

The experiment consisted of three types of trials: no-vision, unperturbed, and perturbed trials (Figure 1A). At the start of each trial, participants held the cursor in the origin. The target appeared after one second, indicating that the participant should start the movement. In all trial types, the cursor disappeared when the handle left the origin. In no-vision trials, the cursor did not reappear during the entire movement. In unperturbed and perturbed trials, the cursor reappeared when the handle distance from the origin exceeded 5cm. However, in perturbed trials, the cursor was projected at a predefined angle relative to the actual handle position.

**Figure 1:**
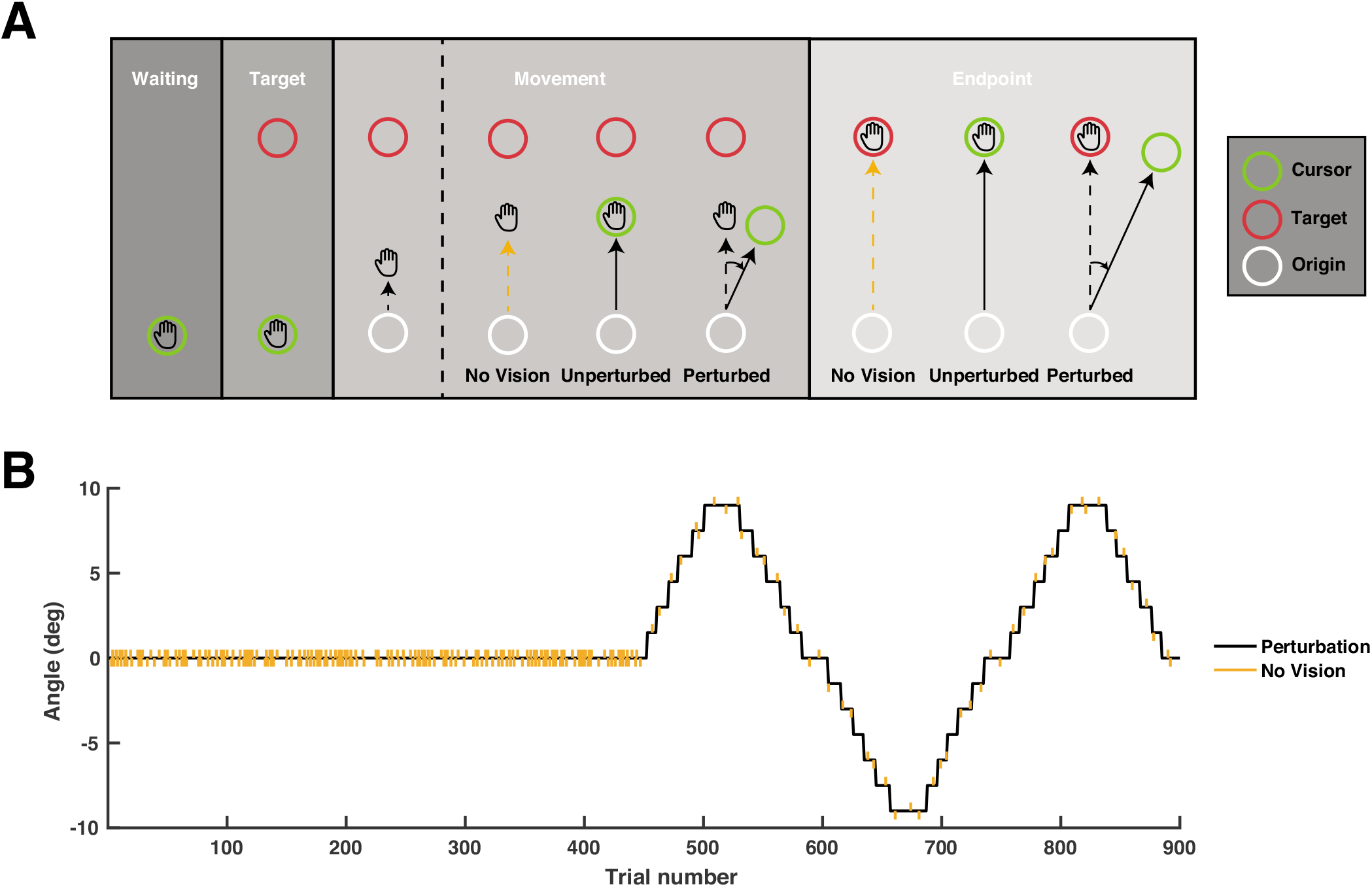
Design of the experiment. **A:** Trial design. **B:** Experimental design.

The movement was damped with a force cushion (3.6Ns/m, ramped up over 7.5ms) when the handle position exceeded 10cm distance from the origin, and thus exceeded the target distance. We defined this time point as movement offset. In perturbed and unperturbed trials, the cursor froze at this time point to provide feedback on the endpoint error. Furthermore, the target changed color. If the movement duration was too short (<400ms), the target stayed red, if the movement duration was correct (400-600ms), the target turned white, and if the duration was too long (>600ms), the target turned blue. We instructed participants that these colors were merely a guideline to ensure ballistic movements without online adjustments and emphasized that the goal of the task was to hit the target. One second after onset of the force cushion, the robot pushed the handle back to the starting position. When the handle was at the origin, the cursor reappeared at the handle position.

The experiment consisted of 900 trials, divided into two blocks of 450 trials: a baseline block followed by a perturbation block (Figure 1B). The exact order of the trial types was randomized for each participant. The baseline block contained 225 unperturbed trials and 225 no-vision trials in a completely randomized order. In contrast, the perturbation block contained 400 perturbation trials and 50 no-vision trials in a pseudorandomized order: in every epoch of 9 trials, there was one randomly interspersed no-vision trial. Thus, the experiment contained 625 trials with visual feedback. In the perturbation block, the perturbation angle changed from 0° to 9° to −9° and back to 0° with increments of 1.5° every 8 to 12 trials. The experiment was paused after every 150 trials for approximately 2 minutes.

### Movement recording and preprocessing

The experiment was controlled by a custom C++ program. The position and velocity signals of the robot handle were sampled at 500Hz. The velocity signal was filtered with an exponential moving average filter (smoothing factor = 0.18s). Movement onset was defined as the time point when movement velocity exceeded 0.03 m/s and movement offset was defined as the moment that the hand location exceeded 10cm distance from the starting position. The hand angle was defined as the angle between the vector connecting the origin and the hand at the end of the movement, relative to the vector connecting the origin and the target. The clockwise direction was defined as positive. Trials were removed if movement duration exceeded 1s or if the absolute hand angle exceeded 30 degrees. On average, 1% (range [0 18]) of the total 900 movement trials were excluded per participant.

### Movement analysis

From the movement recordings, we estimated the execution noise, planning noise, and adaptation rate of each participant. The analysis was performed with Bayesian Markov Chain Monte Carlo simulations in JAGS (Plummer, 2017) and is extensively described in van der Vliet et al. (2018). In short, we fitted a state-space model of trial-to-trial behavior (Cheng and Sabes, 2006).

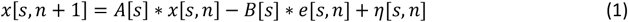

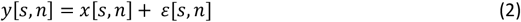

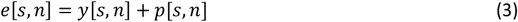

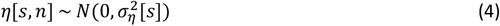

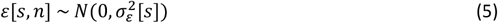

For each trial *n* of participant *s*, *x*[*s, n*] is the movement plan, *y* [*s, n*] the angle of the hand relative to the target at the endpoint, *p*[*s, n*] the perturbation angle and *e*[*s, n*] the error angle of the cursor on the screen relative to the target. Participant-specific motor adaptation parameters estimated with this model are the retention rate *A*[*s*], which is the fractional retention of the movement plan *x*[*s, n*] in the previous trial, and the adaptation rate *B*[*s*], which is the fractional change caused by the error *e*[*s, n*] in the previous trial. The noise terms include planning noise *η*[*s, n*], and execution noise *ε*[*s,n*]. Planning noise and execution noise are modeled as zero-mean Gaussians. The standard deviations of these Gaussians *σ_η_*[*s*] and *σ_ε_*[*s*] represent the magnitude of planning and execution noise for each participant. The participant-specific motor learning parameters were estimated hierarchically with uninformative priors. Additionally, we calculated the lag-1 autocorrelation (R(−1)), and the standard deviation of the signed errors (*σ_y_*) for each participant.

### EEG recording and preprocessing

Participants were wearing a 128 channel EEG cap connected to a 136 channel REFA system (TMSi, Oldenzaal, The Netherlands). The EEG data were recorded with a sampling rate of 2,048Hz from 62 EEG channels: FP1, FPz, FP2, AF7, AF3, AF4, AF8, F7, F5, F3, F1, Fz, F2, F4, F6, F8, FT7 FC5, FC3, FC1, FCz, FC2, FC4, FC6, FT8, T7, C5, C3, C1, Cz, C2, C4, C6, T8, TP7, CP5, CP3, CP1, CPz, CP2, CP4, CP6, TP8, P7, P5, P3, P1, Pz, P2, P4, P6, P8, PO7, PO5, PO3, POz, PO4, PO6, PO8, O1, Oz, O2, and 2 EOG channels. The EOG electrodes were located between the eyebrows and to the right of the right eye. Impedance was kept below 4 kΩ. The recording of each channel was referenced to the average signal across all channels.

After recording, the EEG signals were preprocessed with the EEGLAB toolbox (version 14.1.2b; Swartz Center for Computational Neuroscience, La Jolla, USA) and custom scripts in MATLAB version 2018b (Mathworks, Natick, USA). The raw average referenced EEG signals were digitally filtered between 2-40Hz with a 6^th^ order Butterworth filter, cut into trial epochs of 2000ms centered around the onset of visual feedback, and down-sampled to 128Hz. Non-stereotypical artifacts were automatically removed by excluding trials in which the average log power was an outlier compared to the other trials within the same channel (absolute z-score >3.5). Data quality of the remaining trials was verified by visual inspection. Similarly, stereotypical artifacts were automatically removed by performing a fast-independent component analysis (Delorme et al., 2007) and excluding components in which the average log power was an outlier (absolute z-score >3.5). Visual inspection confirmed that the removed components corresponded to eye movements. The other components were projected back to the channel space. On average, 40 (range [14 79]) of the 900 EEG traces were excluded and 1.1 (range [1 2]) of the 64 components were removed. The remaining traces were used for analysis in the time domain.

For the analysis in the frequency domain, the remaining EEG traces were first filtered with a surface Laplacian to improve the spatial resolution (Perrin et al., 1989). Subsequently, the traces were decomposed into their time-frequency representations using Morlet wavelet convolution applied to the concatenated trial data (Tallon-Baudry et al., 1996; Cohen, 2014). The wavelet frequencies ranged from 2 to 30 Hz in 29 steps of 1Hz (with the number of cycles ranging from 4 to 10 in 29 logarithmically-spaced steps, respectively, for the 29 frequencies). After convolution, the deconcatenated traces were trimmed between –800ms and 800ms, and centered on the onset of visual feedback to remove edge artifacts. Consecutively, the log power at each time-frequency point within each trial was normalized as a percentage change relative to the average log power of that frequency in the predefined baseline window (from 800ms until 600ms before visual feedback) across all trials for the subject.

### EEG analysis

The EEG analysis had three parts: a trial level analysis in the time domain, a trial level analysis in the frequency domain and a participant level analysis. We used linear mixed-effects models (MATLAB’s fitlme function) to estimate trial level effects across participants and a multivariate linear regression (MATLAB’s fitlm function) to estimate effects on the participant level. All variables were z-score normalized for the purpose of the regression. Tables show the normalized effects in order to compare their relative strength. When effect sizes are shown in their original units, they have been transformed back using the standard deviations of the dependent and independent variables.

A signed error was defined as the angle between the cursor on the screen relative to the target; error magnitude was defined as the absolute of the signed error; and error correction was defined as the hand angle in the following trial minus the hand angle in the corresponding trial, multiplied with the sign (1 or −1) of the error angle in the corresponding trial. Thus, negative values represent corrections in the direction of the target, whereas positive values represent corrections in the opposite direction. In the EEG analysis, we only included trials with visual feedback.

For the trial level analysis in the time domain, a separate regression was performed for different time windows. We used a fine-grained collection of windows for generating figures. For a given time window, we regressed EEG amplitude on signed error, error magnitude, and error correction, as well as possible confounding variables such as the number of previous trials and the feedback color (which indicated whether movement duration was within the desired range, see above).

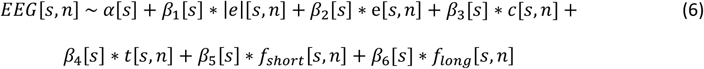

For each trial *n* of participant *s, EEG*[*s, n*] is the average EEG amplitude in a time window, |*e*|[*s, n*] is the movement error magnitude, *e*[*s, n*] is the signed error, *c*[*s, n*] is error correction, *t*[*s, n*] is the original trial number and *f_short_*[*s, n*] and *f_long_*[*s, n*] are binary variables indicating if the movement duration was “too short” (*f_short_*[*s, n*] = 1) or “too long” (*f_long_*[*s, n*] = 1). All participant-specific estimates were modeled as random effects around the group average fixed effects. Throughout the paper, we often refer to the coefficient of the error magnitude (*β*_1_[*s*]) as the EEG-error-sensitivity.

For the trial level analysis in the frequency domain, a separate regression was used for each time-frequency window. We used both fine-grained time-frequency binning to generate figures and a larger bin (4-8Hz and 100-600ms) to analyze frontal midline theta activity in channel FCz (Cavanagh et al., 2012; Arrighi et al., 2016; Savoie et al., 2018). This bin was chosen after visual inspection of the fine-grained picture. For these analyses, we use the linear mixed-effects model in Eq. 6, but with *EEG*[*s, n*] representing the average EEG power in the time-frequency bin being analyzed. We compared different versions of the model with a backward elimination method. Stepwise removal of variables with the smallest effect size was continued as long as the Bayesian Information Criterion (BIC) decreased. The model for which further removal of the smallest effect size would increase the BIC was taken to be the one that best explained the data. FMΘ-error-sensitivity was defined as the relation between frontal midline theta power (FMΘ) and error magnitude in this final model (*β*_1_ [*s*]).

For the participant level analysis, we tested for linear relationships between FMΘ-error-sensitivity (*β*_1_[*s*]), planning noise *σ_η_*[*s*], execution noise *σ_ε_*[*s*], and adaptation rate *B*[*s*]. We tested two regression models specifically: FMΘ-error-sensitivity regressed on the noises and adaptation rate, and adaptation rate regressed on the noises and the FMΘ-error-sensitivity. The choice of these two models reflects an underlying assumption that the noises characterize an individual while adaptation and neural signals vary under different conditions.

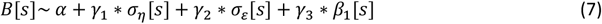

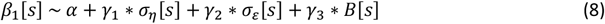

In these models, we used the participant-specific point estimates from the movement analysis and the trial level EEG analysis. In order to compare the strength of the correlations, all variables were z-score normalized. Stepwise removal of variables with the smallest effect size was continued as long as the Bayesian Information Criterion decreased, and the last model for which this criterion held was selected. As an internal control, we also investigated whether EEG sensitivity to the signed error *β*_2_[*s*]) in occipital channels PO5 and PO6 (4-8Hz and 0-500ms) was related to the participant-specific motor learning parameters.

After performing the analysis for the first time, we redid the trial level analysis in the frequency domain with error probability (*β*_7_[*s*]) instead of error magnitude (*β*_1_[*s*]) (Eq. 6). Error probability was defined as the probability for an error magnitude larger than the error magnitude in the corresponding trial, assuming a normal distribution for the errors.

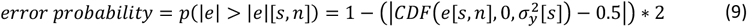

In formula 9, *CDF*(*x,μ,σ*) is the cumulative density function of the normal distribution with mean *μ* and standard deviation *σ* at the point x.

### Data and code availability

The JAGS code for the Bayesian state-space model can be accessed without restrictions at https://github.com/rickvandervliet/Bayesian-state-space. Other scripts and data are available on reasonable request.

## RESULTS

### Movement analysis

On average, target onset, movement onset, and movement offset occurred at −441ms (range across participants [−512 −401]), −193ms [−247 −151] and 78ms [50 118] respectively, relative to the onset of visual feedback. Over the course of the experiment, participants adapted their reaching movements to the perturbation signal (Figure 2). As a result, the average error magnitude was similar in the baseline block and the perturbation block (Figure 2). In 40% [23 58] of the trials, the cursor was completely inside the target (error magnitude < 1.43°) and in an additional 47% [36 51] of the trials, the cursor was partially inside the target (error magnitude < 4.29°) (Figure 3). Figure 3 also highlights the error distribution of 5 participants with different error distributions whose data will be used as example data in subsequent figures. The Bayesian fitting procedure showed that the average planning noise, execution noise, and adaptation rate were 0.47° [0.23 0.78], 2.46° [1.40 4.49], and 0.14 [0.06 0.27], respectively. The movement data of the 5 example participants, ordered by increasing execution noise, is illustrated in Figure 4.

**Figure 2:**
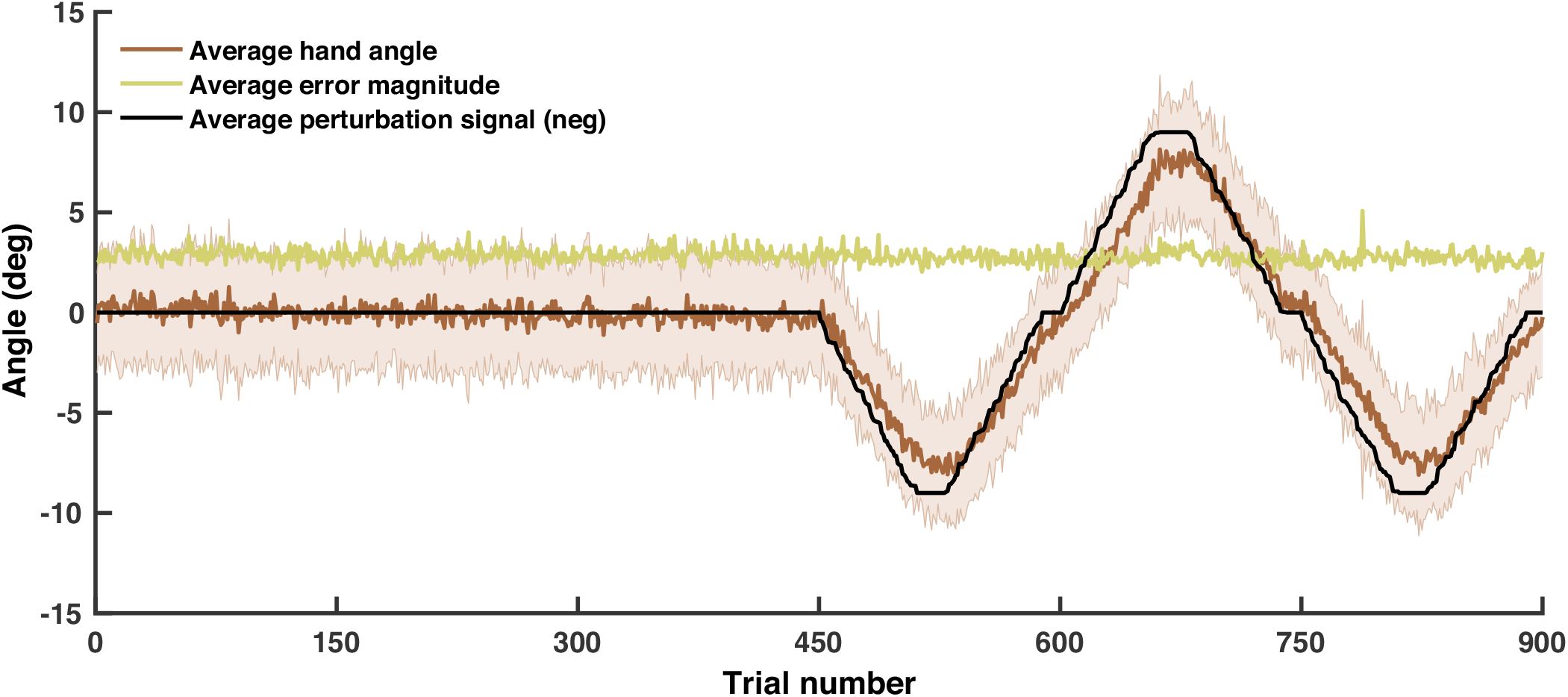
Average movement data across participants throughout the experiment. The error bars on the brown line indicate the standard deviation of the hand angle across subjects.

**Figure 3:**
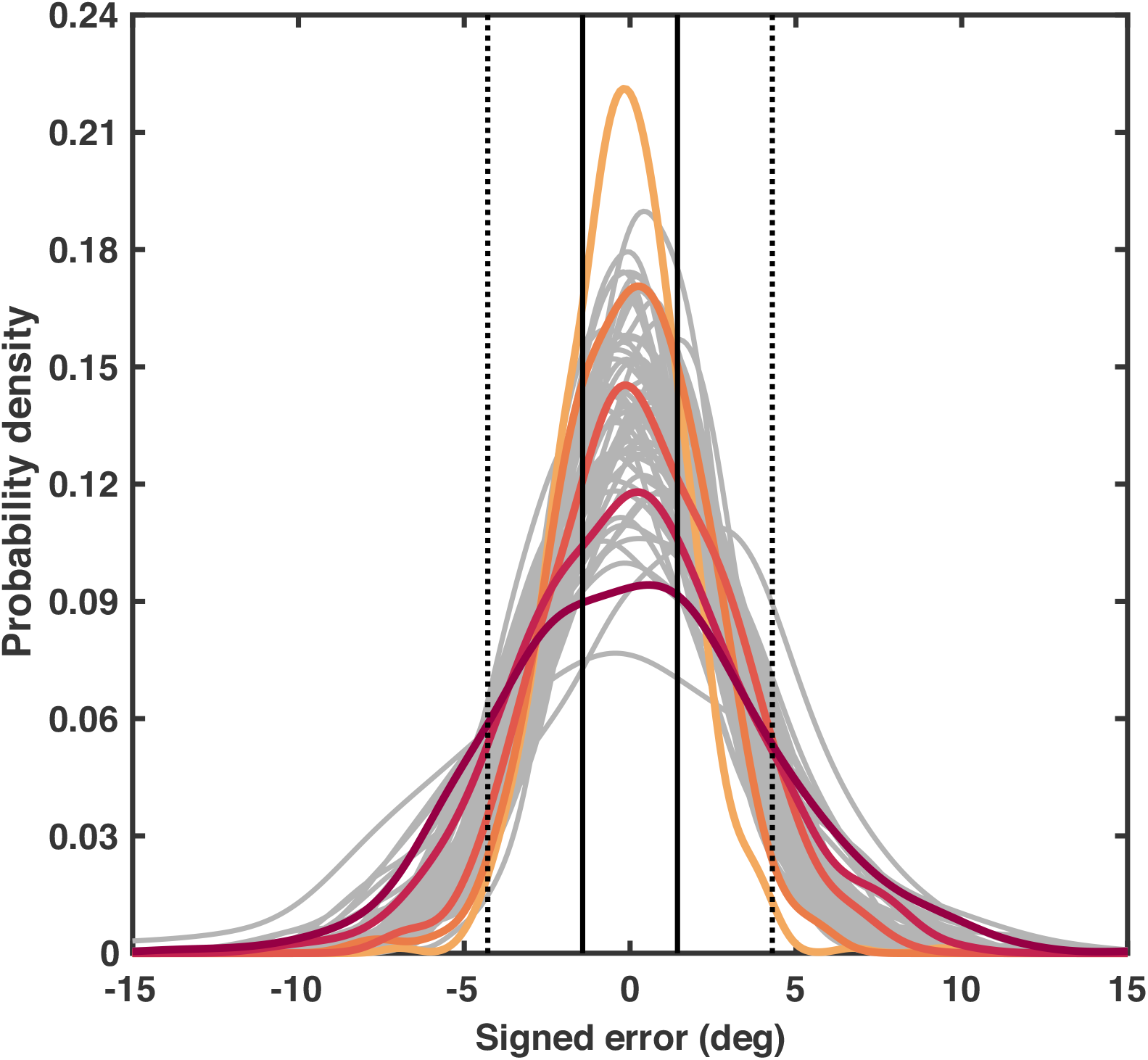
The probability density function of the signed errors for each participant. In trials with error magnitudes below 1.43 degrees (solid black line), the cursor was completely inside the target and in trials with error magnitudes between 1.43 and 4.29 degrees (dotted black line), the cursor was partially in the target.

**Figure 4:**
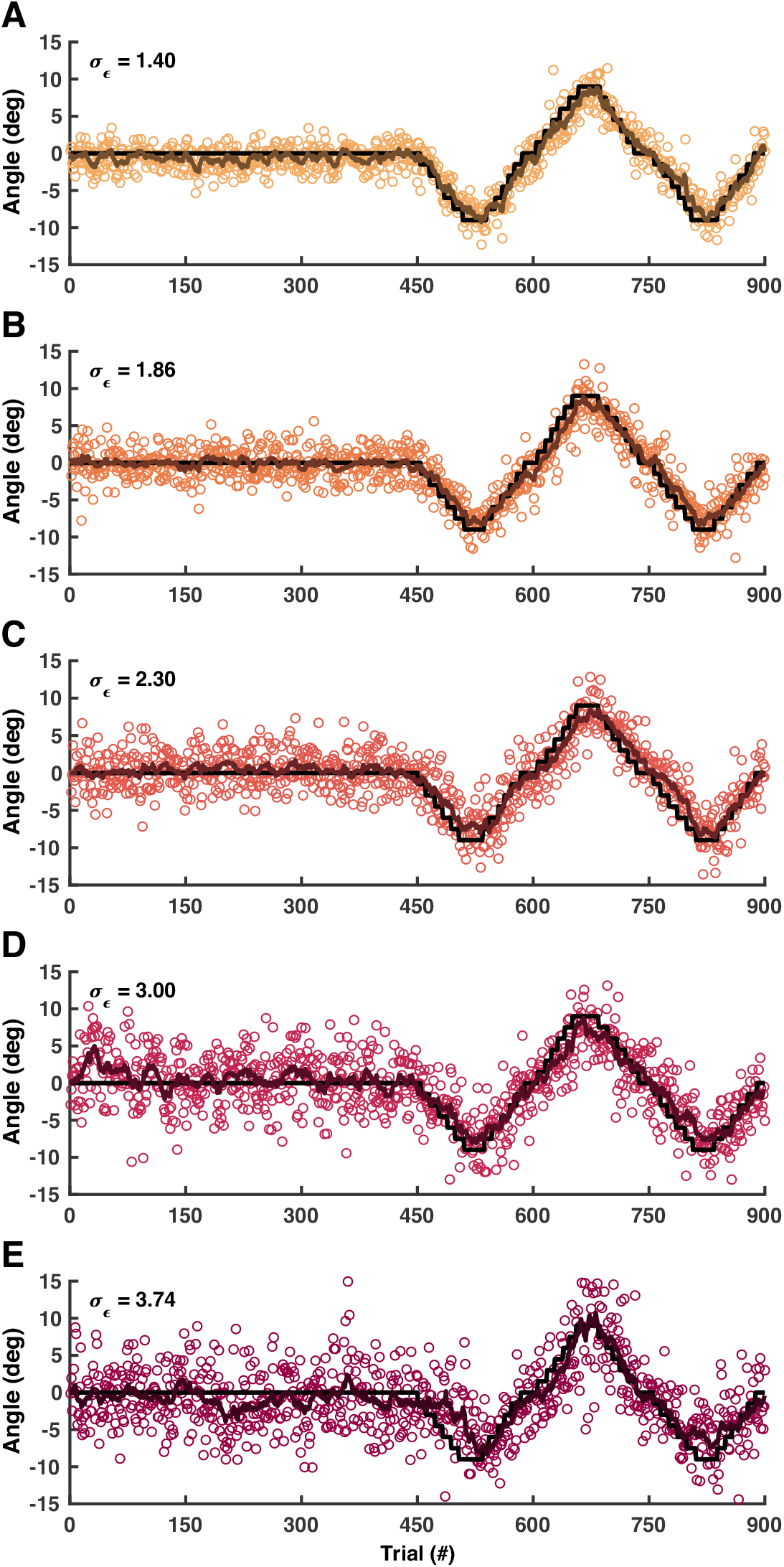
Individual movement data of five individuals with increasing execution noise (*σ_ε_*). Colored circles represent the hand angle in individual trials. The colored lines represent the estimated movement plan from the Bayesian fitting procedure.

We inspected the validity of the Bayesian fitting procedure by correlating the estimated participant-specific factors with the lag-1 autocorrelation, and the standard deviation of the signed errors (Supplementary Figures 1-2). As expected, planning noise was related to the lag-1 autocorrelation in the baseline block (normalized slope = 0.58, CI95% = [0.36 .79]) and execution noise was strongly related to the standard deviation of signed errors throughout the experiment (normalized slope = 0.97, CI95% = [0.91 1.03]).

### EEG analysis

In the time domain, the overall EEG amplitude showed a negative peak over the motor cortices between approximately −100 and 100ms and a large positive peak between approximately 200 and 400ms after visual feedback distributed spatially from the frontal midline region back to the parietal cortex (Figure 5A). Only the positive peak showed a correlation with error magnitude. Within this larger response, the EEG amplitude over the frontal midline region showed negative sensitivity to error magnitude between 200 and 300ms and positive sensitivity between 300 and 400ms (Figure 5B). The correlation with signed error was much smaller, but some relationships could be seen in the occipital cortices between approximately 100 and 300ms (Figure 5C). Any relationship between EEG amplitude and error correction was vanishingly small (Figure 5D).

**Figure 5:**
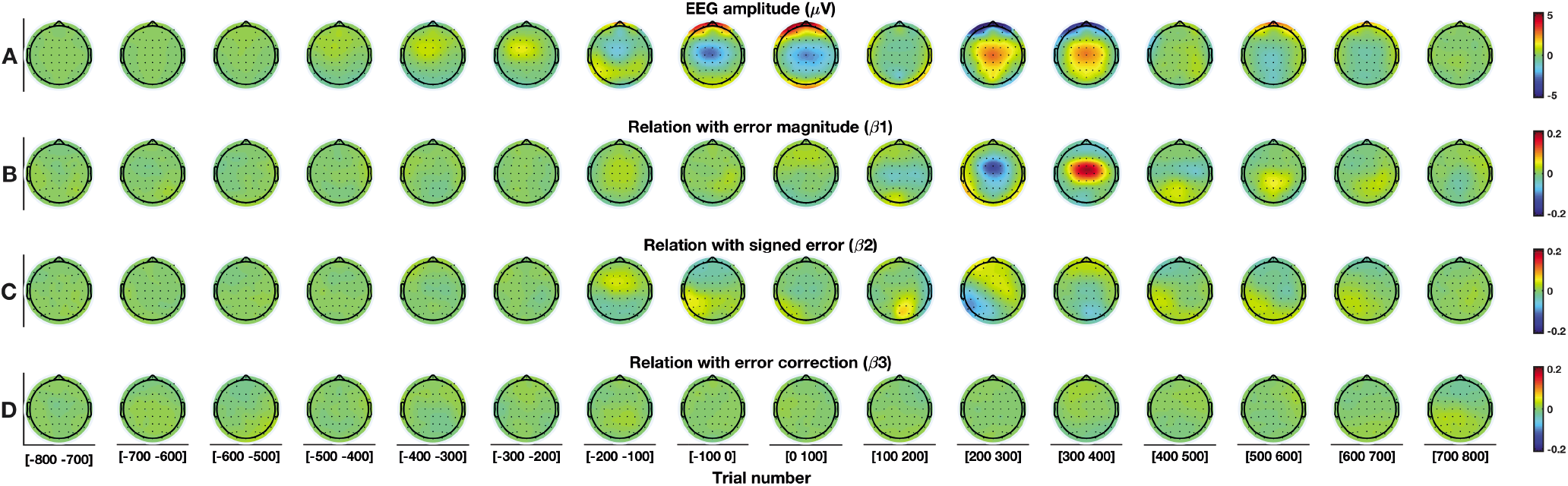
Trial level analysis in the time domain. For this figure, the full mixed-effects model is used (Eq. 6). **A:** EEG amplitude. **B:** Relation with error magnitude. **C:** Relation with signed error. **D:** Relation with error correction.

In the frequency domain, overall theta power showed widespread synchronization between −400ms (approximately target onset) and 600ms (Figure 6A). This can be divided into two phases with a first phase before visual feedback over the frontal midline region and the somatosensory cortices and a second phase after visual feedback over the frontal midline region and the occipital cortices, but then migrating to the prefrontal cortex after approximately 400ms. Within this second synchronization, theta power over the frontal midline region was related to error magnitude (Figure 6B) and also weakly related to signed error in the counterclockwise direction (Figure 6C). Furthermore, theta power in the left and right occipital cortices was related to signed error in the contralateral direction (Figure 6C). Finally, we did not find a relation between theta power and error correction (Figure 6D). Figure 7 illustrates that EEG-error-sensitivity in the frontal midline region (channel FCz) was most prominent in the theta (4-8Hz) frequency band, with foothills in the delta (2-3Hz) and alpha (9-14Hz) frequency bands. These foothills of EEG-error-sensitivity are also visible in the scalp plots of delta and alpha power (Supplementary Figures 3-4). In this experiment we did not see a relation between beta power and error magnitude, signed error, nor error correction (Supplementary Figure 5).

**Figure 6:**
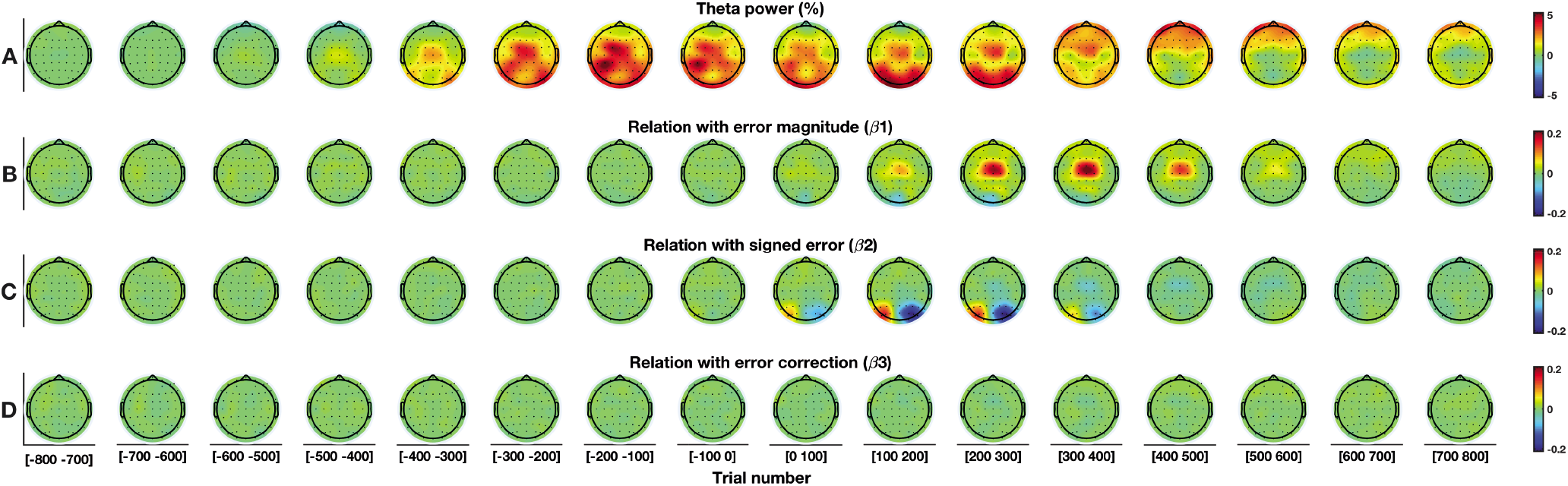
Trial level analysis in the theta frequency band (4-8Hz). For this figure, the full mixed-effects model is used (Eq. 6) **A:** Theta power. **B:** Relation with error magnitude. **C:** Relation with signed error. **D:** Relation with error correction.

**Figure 7:**
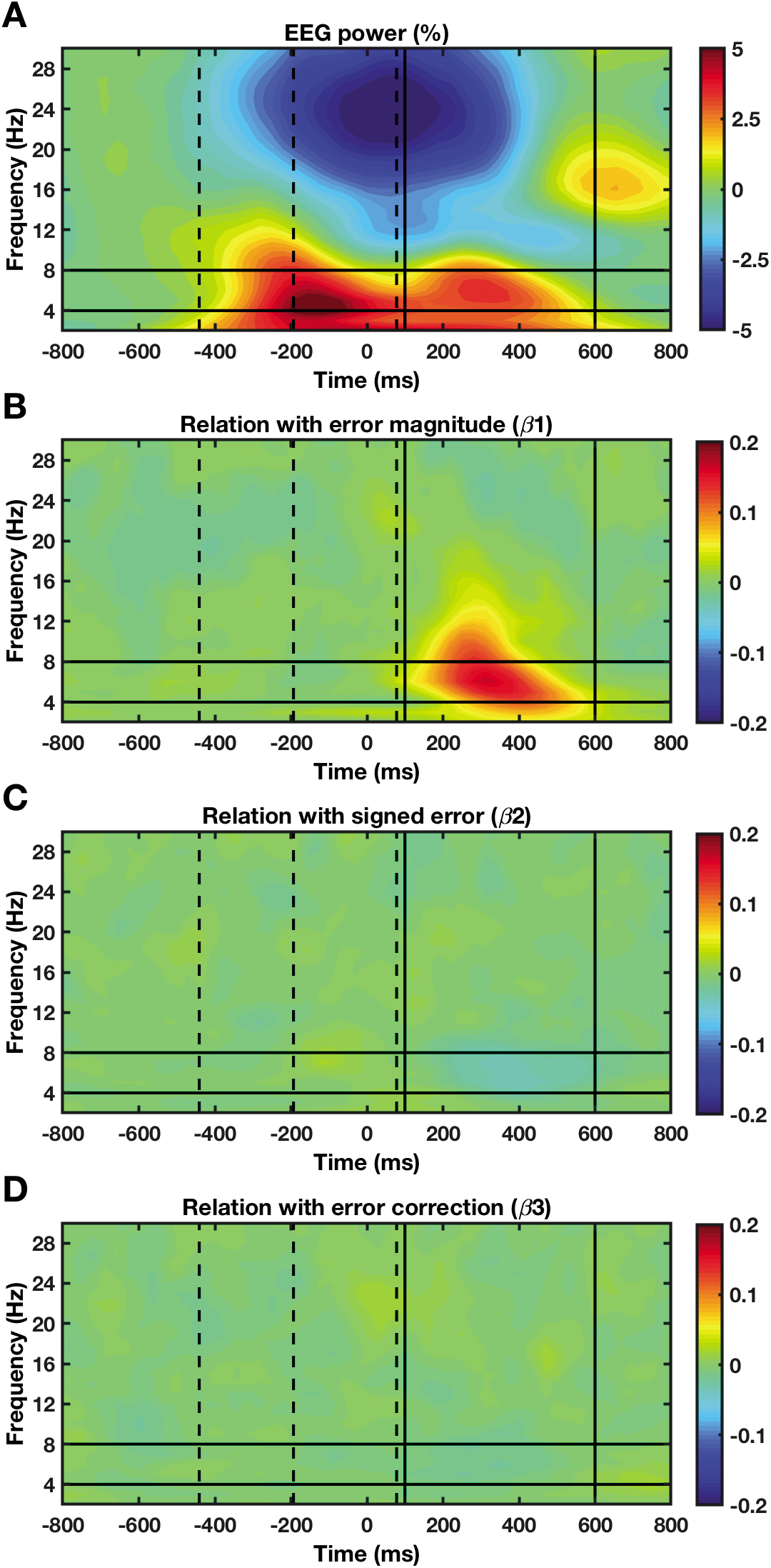
Trial level analysis in the frequency domain in channel FCz. For this figure, the full mixed-effects model is used (Eq. 6). **A:** EEG power. **B:** Relation with error magnitude. **C:** Relation with signed error. **D:** Relation with error correction. Dotted lines represent the average target onset (−441ms), movement onset (−193ms) and movement end (78ms), relative to the onset of visual feedback. Solid lines represent the post-feedback window (100-600ms) and the theta frequency band (4-8Hz).

Figure 8 illustrates that the positive and negative error related peaks over the frontal midline region (Figure 8A-B) are reflections of theta waves in the frequency domain (Figure 8C-D). The signal in the frequency domain contains less noise and is unipolar, which makes the analysis less susceptible for the borders of the timewindow (Supplementary Figure 6). In channel FCz, the relation between theta power and error magnitude was visible from approximately 100 to 600ms. In this post-feedback window, larger error magnitudes resulted in higher FMΘ (Figure 8F). This relation was slightly stronger for errors in the counterclockwise direction (Figure 8E). Backward elimination of regression terms confirmed that post-feedback FMΘ activity was related to error magnitude (FMΘ-error-sensitivity), the number of preceding trials, and the signed error, and not to error correction nor changes in the target color as feedback on the movement duration (Table 1). The average FMΘ-error-sensitivity was 0.57% change in theta power per 1 degree of error magnitude (range across participants [0.07 1.84]). While post-feedback FMΘ activity gradually decreased over the course of the experiment, FMΘ-error-sensitivity was consistent across the baseline and perturbation blocks (Figure 8G). Furthermore, Supplementary Figure 7 illustrates that FMΘ-error sensitivity was also present within the subset of trials that were completely inside the target (that is when error magnitude < 1.43).

**Figure 8:**
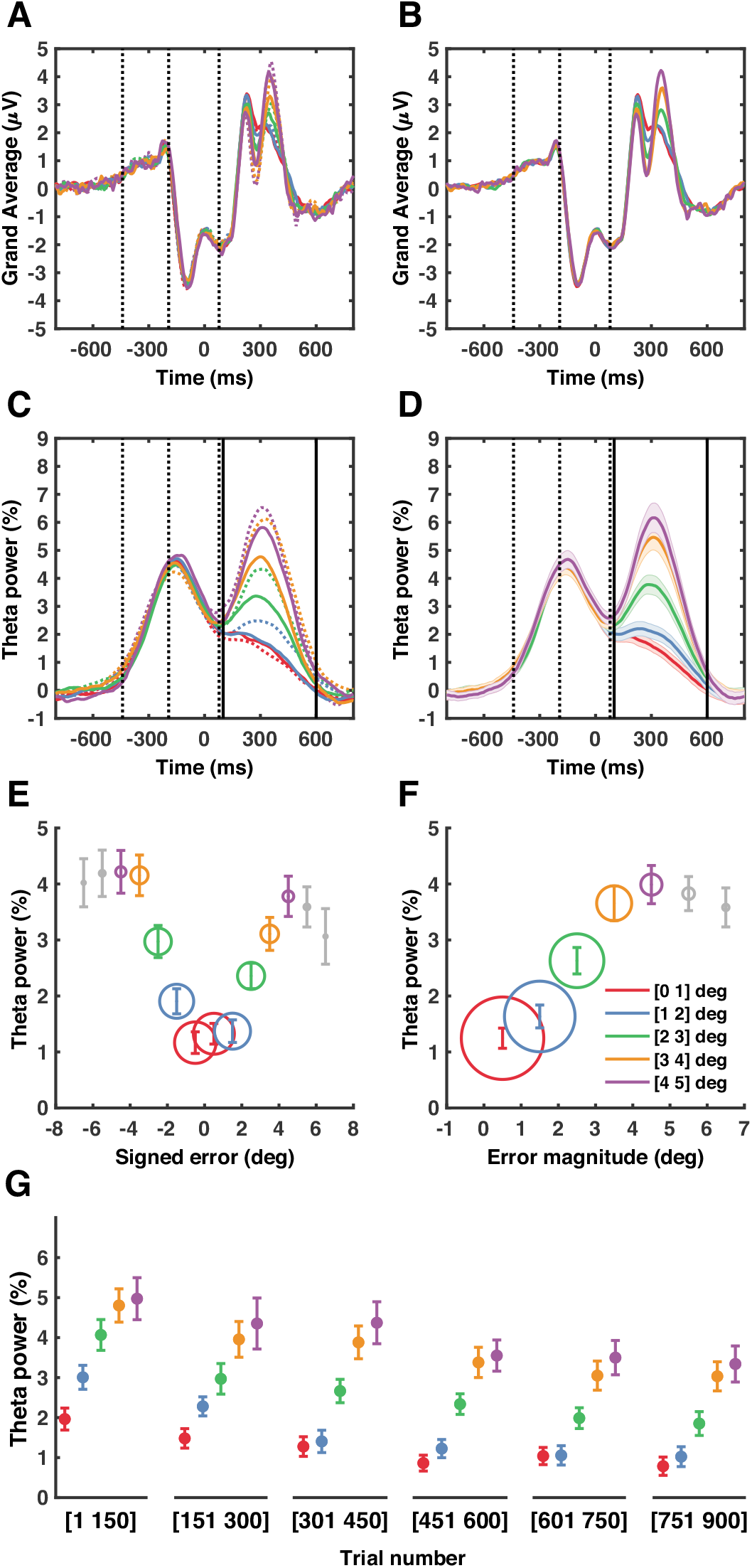
Error sensitivity in channel FCz. For visualization purposes, trials have been binned and averaged based on signed error (**A,C,E**) or error magnitude (**B,D,F,G**). Colored dotted lines represent bins with a negative sign. Black dotted lines represent the average target onset (−441ms), movement onset (−193ms) and movement end (78ms), relative to the onset of visual feedback. Black solid lines represent the post-feedback window (100-600ms). **A-B:** Average EEG amplitude across subjects. **C-D:** Average theta power (4-8Hz) across subjects. In panel D, error bars represent the standard error of the mean. **E-F:** Average theta power in the in the post-feedback window. The diameter of the circles is related to the average number of trials in the bin. **G:** Average theta power in the post-feedback window throughout the experiment. Trials 1-450 are part of the baseline block and thus without any perturbations.

**Table 1:**
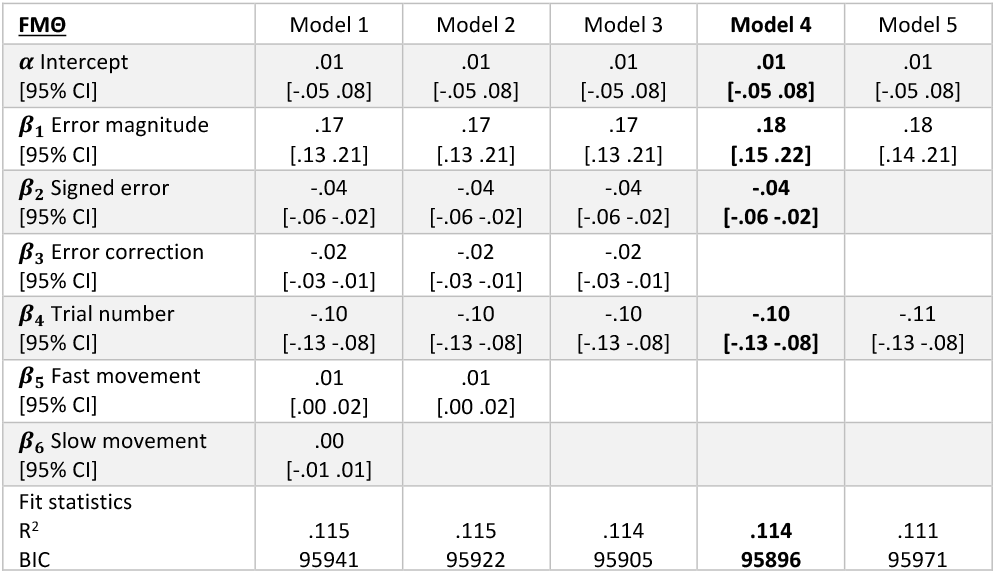
Linear mixed-effects model on the trial level (35117 trials, 60 participants) with frontal midline theta power (FMΘ) as the dependent variable (Eq. 6). In order to compare the strength of the correlations, all variables were z-score normalized. Stepwise removal of variables with the smallest effect size was continued as long as the Bayesian Information Criterion (BIC) decreased, and the last model for which this criterion held was selected. Trial-to-trial differences in FMΘ were best explained by error magnitude, the number of preceding trials and the signed error.

On the participant level, individual differences in adaptation rate were positively correlated to planning noise and negatively related to execution noise and were not further explained by FMΘ-error-sensitivity (Table 2, Figure 9). Conversely, individual differences in FMΘ-error-sensitivity were best explained by execution noise (Table 3, Figure 10). The control analysis showed that individual differences in the occipital sensitivity to signed error (Supplementary Table 1, Supplementary Figure 8) were not related to execution noise (Supplementary Table 2-3, Supplementary Figure 9).

**Figure 9:**
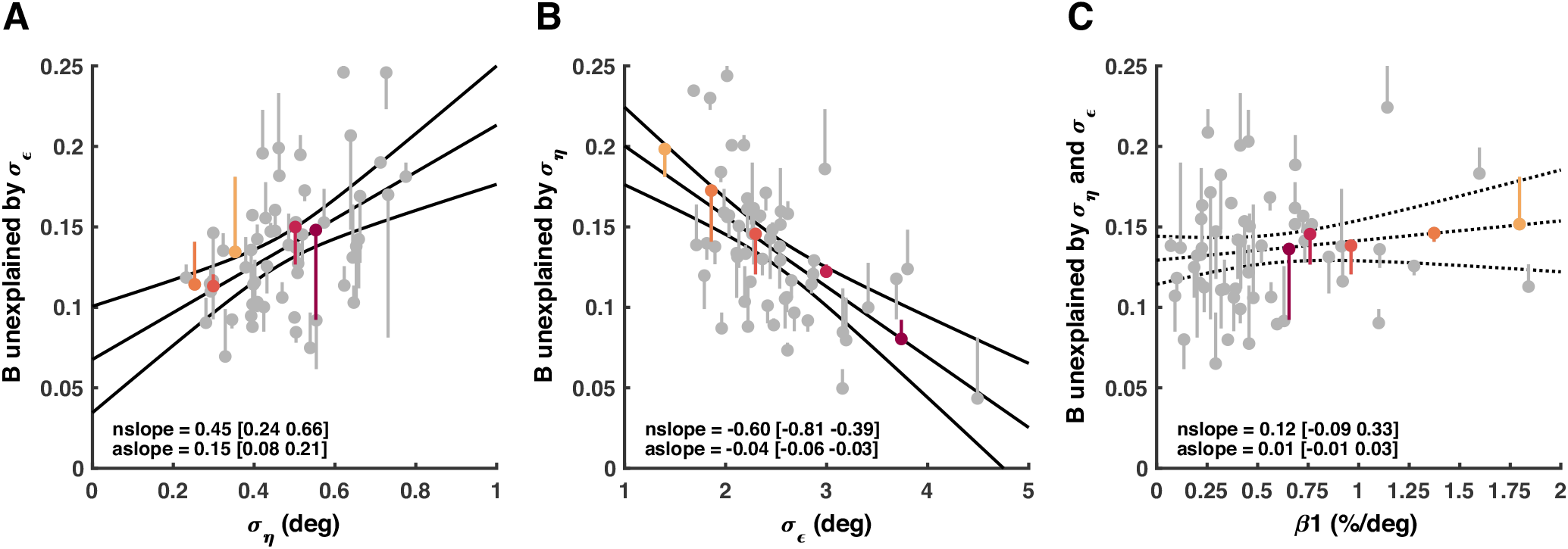
Linear model on the participant level (60 participants), with adaptation rate (B) as the dependent variable (Eq. 7). The best model included planning noise (09) and execution noise (*σ_ε_*), but not FMΘ-error-sensitivity (ft) (Table 2, Model 2). The vertical lines represent the dependent variable. The dots at the end of the lines represent the residuals of the dependent variable after subtracting the estimated effect of the (other) independent variables included in the best model. Black lines represent the absolute slope (‘aslope’) and the confidence interval. Note that the confidence intervals of z-score normalized slope (‘nslope’) of planning noise (09) and execution noise (*σ_ε_*) are slightly smaller than in Table 2, because, in this figure, the effect of the other parameter is ‘fixed’ in order to create the residuals.

**Figure 10:**
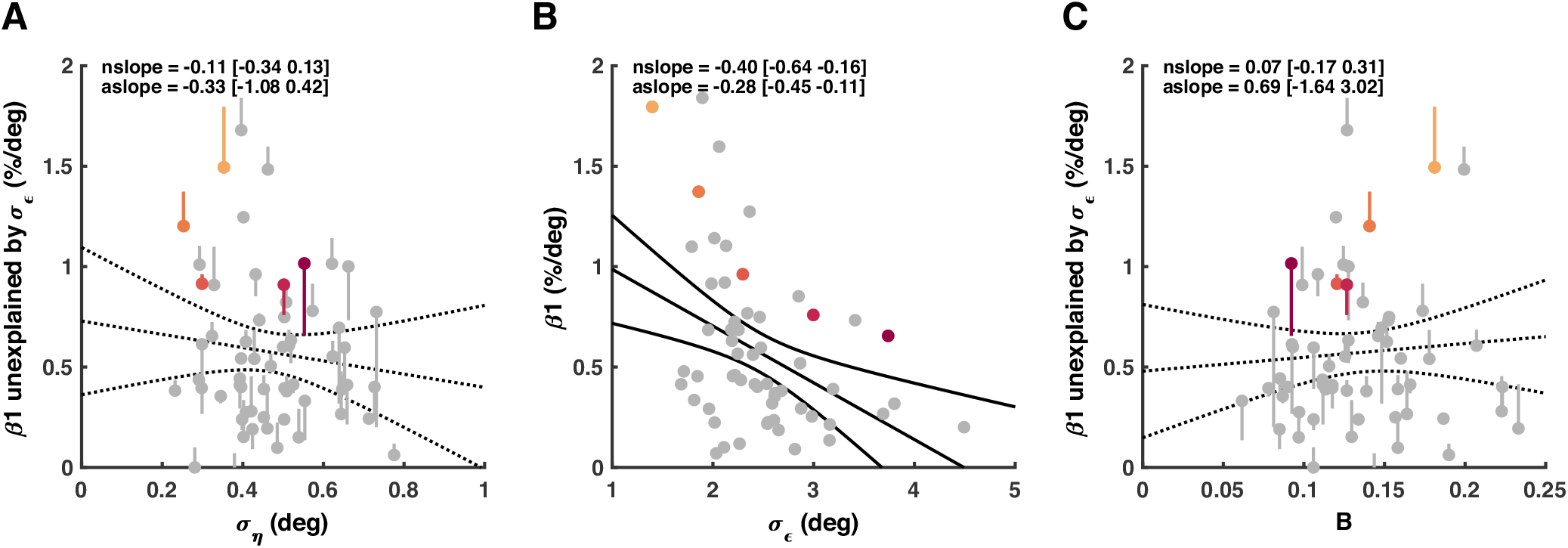
Linear model on the participant level (60 participants), with FMΘ-error-sensitivity (*β*_1_) as the dependent variable (Eq. 8). The best model included execution noise (*σ_ε_*), but not planning (09) noise nor adaptation rate (B) (Table 3, Model 3). The vertical lines represent the dependent variable. The dots at the end of the vertical lines represent the residuals of the dependent variable after subtracting the estimated effect of the independent variables included in the best model. Black lines represent the absolute slope (‘aslope’) and the confidence interval. The z-score normalized slope is also shown (‘nslope’).

**Table 2:**
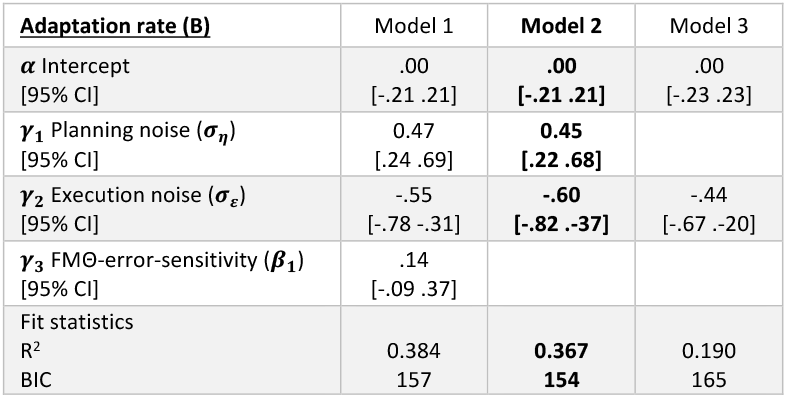
Multivariate linear model on the participant level (60 participants), with adaptation rate (B) as the dependent variable (Eq. 7). FMΘ-error-sensitivity is the relation between frontal midline theta power (FMΘ) and error magnitude on the trial level estimated by the selected model in Table 1. In order to compare the strength of the correlations, all variables were z-score normalized. Stepwise removal of variables with the smallest effect size was continued as long as the Bayesian Information Criterion decreased, and the last model for which this criterion held was selected. Individual differences in adaptation rate were best explained by planning noise and execution noise.

**Table 3:** Multivariate linear model on the participant level (60 participants), with FMΔ-error-sensitivity as the dependent variable (Methods, Eq. 8). FMΘ-error-sensitivity is the relation between frontal midline theta power (FMΘ) and error magnitude on the trial level estimated by the selected model in Table 1. In order to compare the strength of the correlations, all variables were z-score normalized. Stepwise removal of variables with the smallest effect size was continued as long as the Bayesian Information Criterion decreased, and the last model for which this criterion held was selected. Individual differences in FMΘ-error-sensitivity were best explained by execution noise.

| **FMO-error-sensitivity ($\beta_1$)** | Model 1 | Model 2 | **Model 3** | Model 4 |
| --- | --- | --- | --- | --- |
| $\alpha$ Intercept<br>[95% CI] | .00<br>[-.24 .24] | .00<br>[-.24 .24] | **.00**<br>**[-.24 .24]** | .00<br>[-.26 .26] |
| $\gamma_1$ Planning noise ($\sigma_\eta$)<br>[95% CI] | -.20<br>[-.50 .09] | -.12<br>[-.38 .14] | | |
| $\gamma_2$ Execution noise ($\sigma_\epsilon$)<br>[95% CI] | -.24<br>[-.56 .07] | -.35<br>[-.61 -.10] | **-.40**<br>**[-.64 -.16]** | |
| $\gamma_3$ Adaptation rate (B)<br>[95% CI] | .19<br>[-.12 .49] | | | |
| Fit statistics |  |  |  |  |
| R<sup>2</sup> | .193 | .171 | **.158** |  |
| BIC | 173 | 170 | **167** | 173 |

The effect of execution noise level on FMΘ sensitivity led us to consider the possibility that FMΘ reflected the extent to which a given error was expected, considering the individual standard deviation of the signed errors. To test this, we redid the analysis in the frequency domain with error probability instead of error magnitude (Figure 11) and found that this improved the model (Table 4). Using FMΘ sensitivity to error probability instead of error magnitude reduced the individual differences in EEG-sensitivity and effectively removed the relation between EEG-error-sensitivity and execution noise (Table 5, Figure 12). This is further illustrated by the looking at the FMΘ power of the 5 participants who have been used as examples in previous figures (Figure 13).

**Figure 11:**
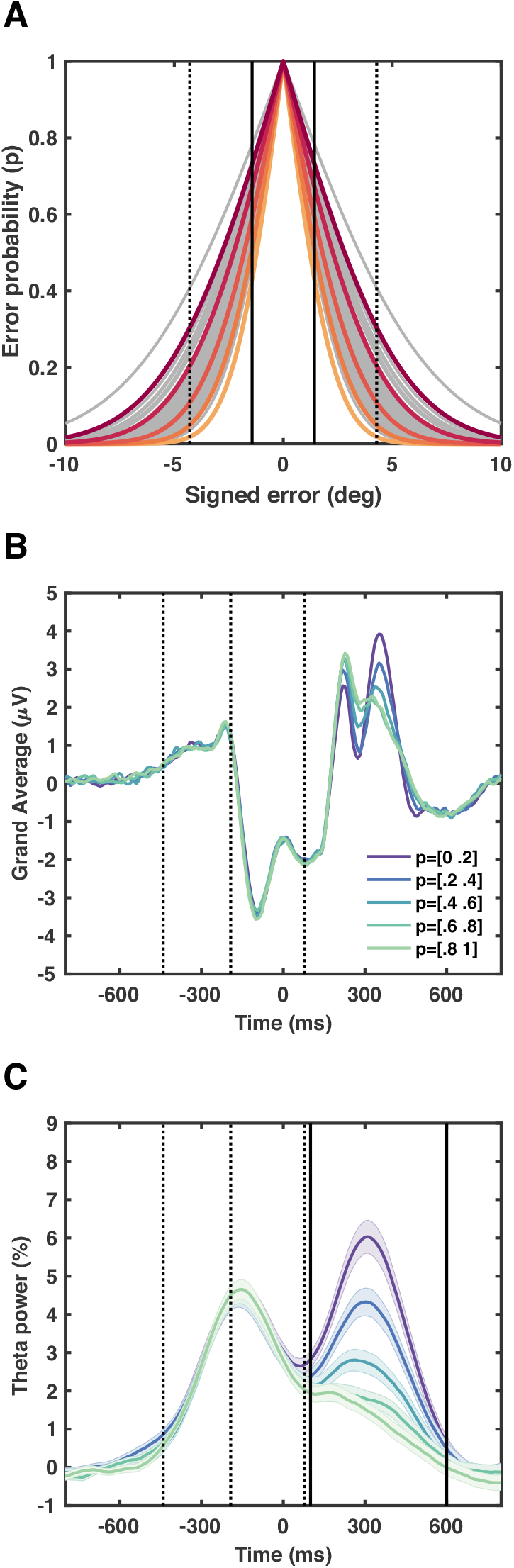
Error probability sensitivity in channel FCz. **A:** Relation between the signed error and error probability for each participant (Formula 9). In trials with error magnitudes below 1.43 degrees (solid black line) the cursor was completely inside the target, and in trials with error magnitudes between 1.43 and 4.29 degrees (dotted black line), the cursor was partially in the target. **B:** Average EEG amplitude across subjects. For visualization purposes, trials have been binned and averaged based on error probability. Dotted lines represent the average target onset (−441ms), movement onset (−193ms) and movement end (78ms), relative to the onset of visual feedback. Solid lines represent the post-feedback window (100-600ms). **C:** Average theta power (4-8Hz) across subjects. Error bars represent the standard error of the mean.

**Figure 12:**
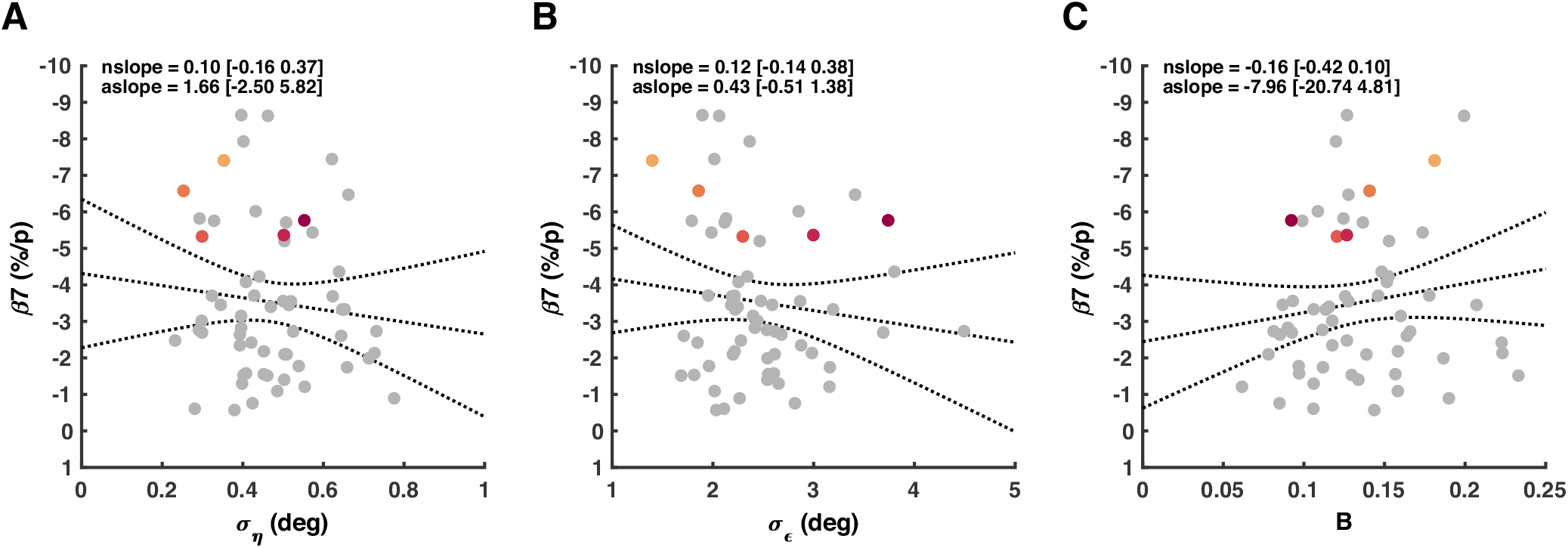
Linear model on the participant level (60 participants), with FMΘ-probability-sensitivity (*β*_7_) as the dependent variable. FMΘ-surprise-sensitivity was not related to planning (*σ_η_*), execution noise (*σ_ε_*), nor adaptation rate (B) (Table 5, Model 4). Black lines represent the absolute slope (‘aslope’) and the confidence interval. The z-score normalized slope is also shown (‘nslope’). Note that the y-axis is inverted for easier comparison with figure 10.

**Figure 13:**
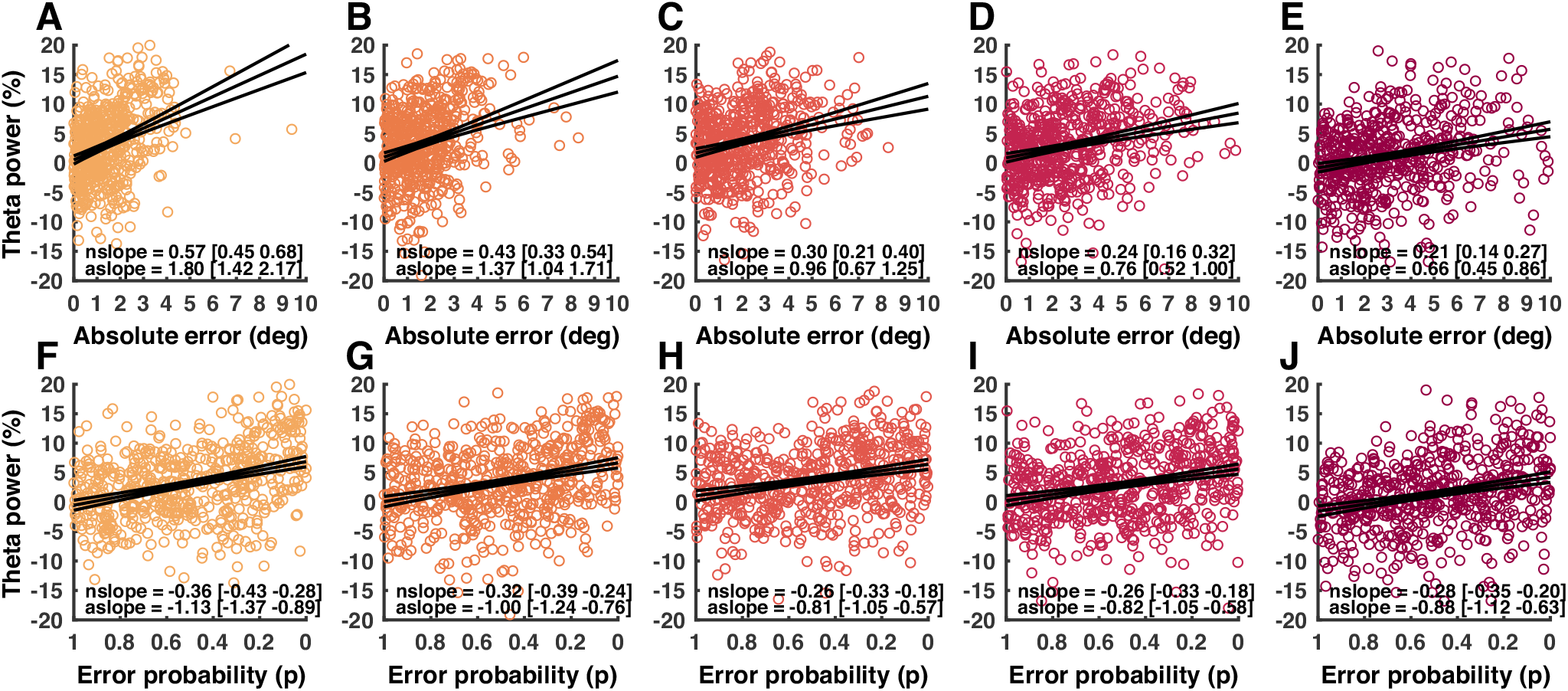
Post-feedback error sensitivity and surprise sensitivity in channel FCz of five individuals with increasing execution noise. Colored circles represent the post-feedback theta power in individual trials. Black lines represent the absolute slope (‘aslope’) and the confidence interval. The z-score normalized slope is also shown (‘nslope’). **A-E**: FMΘ-error-sensitivity. **F-J:** FMΘ-probability-sensitivity. Note that the x-axis is inverted for easier comparison to panel A-E.

**Table 4:** Linear mixed-effects model on the trial level (35117 trials, 60 participants) with frontal midline theta power (FMΘ) as the dependent variable. This table shows the selected model of Table 1, but with error magnitude (*β*_1_) replaced by error probability (*β*_7_). Replacing error magnitude by error probability lowered the Bayesian Information Criterion (BIC).

| **FMO** | **Model 6** |
| --- | --- |
| $\alpha$ Intercept<br>[95% CI] | **.00**<br>[**-.06 .06**] |
| $\beta_7$ Error Probability<br>[95% CI] | **-.17**<br>[**-.20 -.14**] |
| $\beta_2$ Signed Error<br>[95% CI] | **-.04**<br>[**-.06 -.02**] |
| $\beta_4$ Trial Number<br>[95% CI] | **-.11**<br>[**-.13 -.08**] |
| Fit statistics |  |
| R<sup>2</sup> | **.111** |
| BIC | **95730** |

**Table 5:**
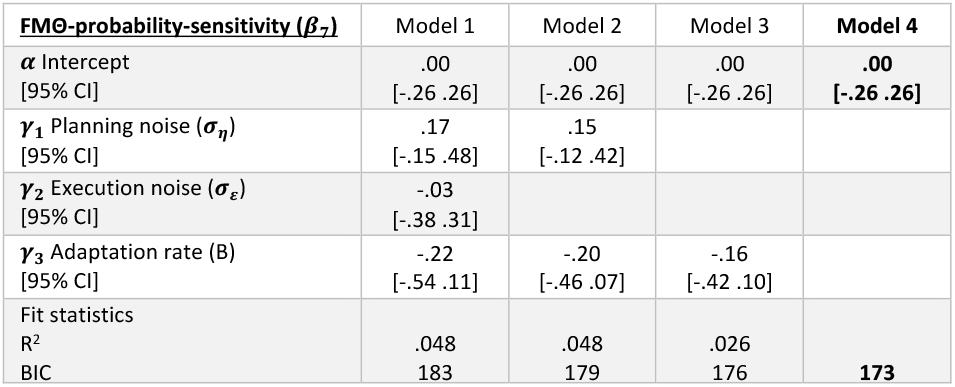
Multivariate linear model on the participant level (60 participants), with FMΘ-probability-sensitivity as the dependent variable. FMΘ-probability-sensitivity is the relation between frontal midline theta power (FMΘ) and error probability on the trial level estimated by the model in Table 4. In order to compare the strength of the correlations, all variables were z-score normalized. Stepwise removal of variables with the smallest effect size was continued as long as the Bayesian Information Criterion decreased, and the last model for which this criterion held was selected. Individual differences in FMΘ-probability-sensitivity were not explained by planning noise, execution noise and adaptation rate.

## DISCUSSION

This study shows that frontal midline theta activity (FMΘ) is also present during very small perturbations, does not drive between-trial error correction and that the sensitivity of FMΘ to error magnitude was smaller for participants with greater execution noise. With 60 participants and, on average, 585 trials (range [430 611]) per participant, this is the largest study on visuomotor adaptation and EEG activity to date, allowing us to investigate individual differences in motor learning and FMΘ. In group average EEG activity, FMΘ appeared to reflect error magnitude (Figure 8). However, this relationship between FMΘ and error magnitude (FMΘ-error-sensitivity) was much weaker in individuals with a larger execution noise and thus a larger standard deviation of signed errors. We reasoned that if FMΘ is affecting subsequent errors, we should see a relation between trial-to-trial differences in FMΘ and error correction. However, our results show no such relationship (Figure 7). Furthermore, we see that individual differences in FMΘ-error-sensitivity are strongly related to execution noise and not to adaptation rate (Figure 10). This left us with an alternative interpretation: FMΘ may reflect a surprise-like saliency signal. We tested this by calculating the error probability in each trial (Figure 11). Defining FMΘ-probability-sensitivity instead of FMΘ-error-sensitivity reduced the individual differences and substantially weakened the correlation with execution noise (Figure 13). These findings are consistent with the hypothesis that FMΘ reflects surprise.

This study replicates our earlier findings that individual differences in adaptation rate are positively related to planning noise and negatively related to execution noise (van der Vliet et al., 2018) which strengthens the evidence that the adaptation rate is tuned to the noise terms according to Kalman filter theory (Kalman, 1960). It also expands the previous literature on post-feedback frontal midline activity in three essential ways.

First of all, this study shows that frontal midline theta activity (FMΘ) is present during very small gradual perturbations. Arrighi et al. (2016) only found a relationship between frontal midline activity and error magnitude for error magnitudes above a certain threshold. Furthermore, Palidis et al. (2019) only used very small gradual perturbations, but did not see a relationship between frontal midline activity and error magnitude. Finally, Savoie et al. (2018) found that FMΘ, corrected for error magnitude, was higher when participants were made aware of a perturbation paradigm and instructed how to counteract it. The results of these previous studies suggest that the relation between frontal midline activity and error magnitude may be dependent on awareness of the perturbation paradigm. However, the current study shows that FMΘ-error-sensitivity is present during very small perturbations and even in the absence of perturbations (during movement calibration in the baseline block), making it safe to assume that participants were unaware of external perturbations and that adaptation was implicit.

Secondly, this study shows that trial-to-trial differences frontal midline theta activity (FMΘ) are not related to between trial error corrections. To our knowledge, this is the first study to investigate the relation between EEG activity and adaptation on the trial level and our results are a strong argument that frontal midline activity does not boost implicit motor adaptation, which is thought to be dependent on the olivocerebellar system (Shadmehr and Krakauer, 2008; De Zeeuw et al., 2011).

Third and finally, this study shows that FMΘ is better explained by error probability than by error magnitude. This supports the hypothesis of Torrecillos et al. (2014) that frontal midline theta activity during motor adaptation may represent a surprise-like saliency signal and is in apparent contrast with the interpretation of Palidis et al. (2019) who concluded that frontal midline activity is related to reward error. Interestingly, the difference between our interpretation and theirs echoes a long-standing dispute on frontal midline theta in the cognitive literature (Cavanagh and Frank, 2014; Sambrook and Goslin, 2015). The argument of Palidis et al. was made by process of elimination. In a first experiment with endpoint error feedback, they ruled out a relation between frontal midline activity and the magnitude of the endpoint errors, which were all inside the target (success). In a second experiment with probabilistic reward feedback, they showed a relation between frontal midline activity and reward. Combining these two, they concluded that frontal midline activity in the second experiment was solely related to reward error and not to saliency.

We argue that they may not have found a relation between frontal midline activity and the endpoint errors in their first experiment due to two major differences in their methodology compared to ours. This would weaken the process of elimination argument such that the frontal midline activity in their second experiment may also reflect a surprise-like saliency signal. The first factor is that their analysis was in the time domain and not the frequency domain. The average EEG amplitude of theta oscillations (4-8Hz) in a time window of 150ms may include both positive and negative peaks that partially cancel each other out (Supplementary Figure 6A-B). The second methodological factor is the difference in the statistical power of the two data sets. With fewer participants and fewer trials, the presence and significance of these smaller peaks within a larger positive response may not have been apparent (Supplementary Figure 6C-D). Indeed, our study shows FMΘ sensitivity to error magnitude even in those trials where the cursor was completely inside the target (Supplementary Figure 7).

Nevertheless, an important limitation of our own study is that we cannot exclude the possibility that the participants constructed their own subgoals. FMΘ was not related to feedback on the movement duration, which confirms that participants understood the instructions that the goal of the task was to hit the target. However, participants may have constructed their own subgoals in which endpoints in the middle of the target were more rewarding than endpoints slightly off-center but still completely inside the target. A dedicated experiment which successfully dissociates error and reward signals, possibly using varying target sizes and reward contingencies, is necessary to resolve this important point.

Other limitations are introduced by the focus on post-feedback EEG activity and implicit adaptation. While we find that frontal midline theta does not drive implicit adaptation, the focus on implicit adaptation means that we cannot exclude the possibility that it drives explicit strategy. Another important point is that we do not see error sensitivity in the beta band. This could be due to a trial structure that was not properly designed to uncover beta band error sensitivity. Previously, beta sensitivity to error magnitude has been found in the pre-movement period and the postmovement beta rebound (Tan et al., 2014; Torrecillos et al., 2015). However, we did not have a preparatory pre-movement period in our task. Furthermore, the use of a force cushion and the automatic pushback at the end of each trial may have obscured the post-movement period.

Regarding the relation between frontal midline theta and error probability, we hypothesize that FMΘ may drive cognitive control (Cavanagh and Frank, 2014). While this study convincingly shows that FMΘ is not dependent on awareness, it is still true that FMΘ may influence awareness and cognitive control. In a visuomotor adaptation task with larger errors or otherwise more salient errors, FMΘ power may facilitate awareness of changes in the environment. This, in turn, might evoke cognitive strategies such as cautious movement, memorization or re-aiming. In individuals with a larger execution noise, the same error magnitude may not indicate a change in the external environment and would thus warrant less cognitive control. An alternative to this hypothesis is that FMΘ power reflects estimation uncertainty (Tan et al., 2016). When confronted with unlikely events, individuals may drop the priors of their internal model and exhibit increased search behavior (planning noise) in order to form new priors. Thus, an experiment where estimation uncertainty and the ability to develop explicit strategies are manipulated independently would be an important step in understanding how the FMΘ response ultimately affects behavior.

## ACKNOWLEDGEMENTS

We would like to thank Nicole Malfait and Dimitrios J. Palidis for constructive feedback on the manuscript.

## SUPPLEMENTARY TABLES

**Supplementary Table 1:**
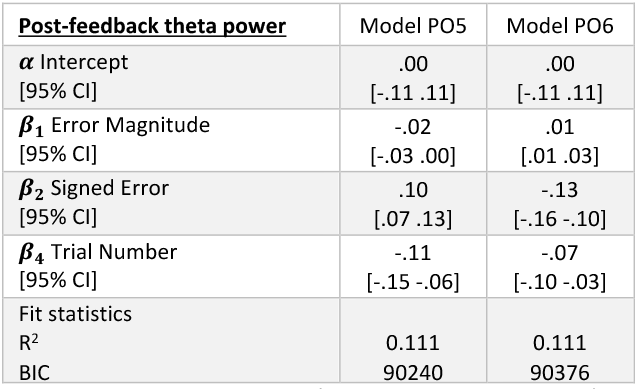
Linear mixed-effects model on the trial level (35117 trials, 60 participants) with theta power (4-8Hz) in the left (PO5) and right (PO6) occipital cortex as the dependent variable. This table shows the selected model of Table 1, but with frontal midline theta power replaced by theta power in the occipital cortices. BIC is the Bayesian Information Criterion.

| **Post-feedback theta power** | Model PO5 | Model PO6 |
| --- | --- | --- |
| $\alpha$ Intercept<br>[95% CI] | .00<br>[-.11 .11] | .00<br>[-.11 .11] |
| $\beta_1$ Error Magnitude<br>[95% CI] | -.02<br>[-.03 .00] | .01<br>[.01 .03] |
| $\beta_2$ Signed Error<br>[95% CI] | .10<br>[.07 .13] | -.13<br>[-.16 -.10] |
| $\beta_4$ Trial Number<br>[95% CI] | -.11<br>[-.15 -.06] | -.07<br>[-.10 -.03] |
| Fit statistics |  |  |
| R<sup>2</sup> | 0.111 | 0.111 |
| BIC | 90240 | 90376 |

**Supplementary Table 2:** Multivariate linear model on the participant level (60 participants), with left occipital (PO5) theta sensitivity to the signed error as the independent variable (Supplementary Table 1, Model PO5). In order to compare the strength of the correlations, all variables were z-score normalized. Stepwise removal of variables with the smallest effect size was continued as long as the Bayesian Information Criterion decreased, and the last model for which this criterion held was selected. Individual differences in theta sensitivity to the signed error were not explained by planning noise, execution noise and adaptation rate.

| **PO5 Theta sensitivity to signed error ($\beta_2$)** | Model 1 | Model 2 | Model 3 | **Model 4** |
| --- | --- | --- | --- | --- |
| $\alpha$ Intercept<br>[95% CI] | .00<br>[-.26 .26] | .00<br>[-.26 .26] | .00<br>[-.26 .26] | **.00**<br>**[-.26 .26]** |
| $\gamma_1$ Planning noise ($\sigma_\eta$)<br>[95% CI] | -.16<br>[-.48 .16] | -.11<br>[-.39 .18] | -.09<br>[-.35 .18] | |
| $\gamma_2$ Execution noise ($\sigma_\varepsilon$)<br>[95% CI] | .13<br>[-.22 .48] | .06<br>[-.22 .34] | | |
| $\gamma_3$ Adaptation rate (B)<br>[95% CI] | .11<br>[-.22 .45] | | | |
| Fit statistics |  |  |  |  |
| R<sup>2</sup> | .019 | -.010 | -.007 |  |
| BIC | 185 | 181 | 177 | **173** |

**Supplementary Table 2:**
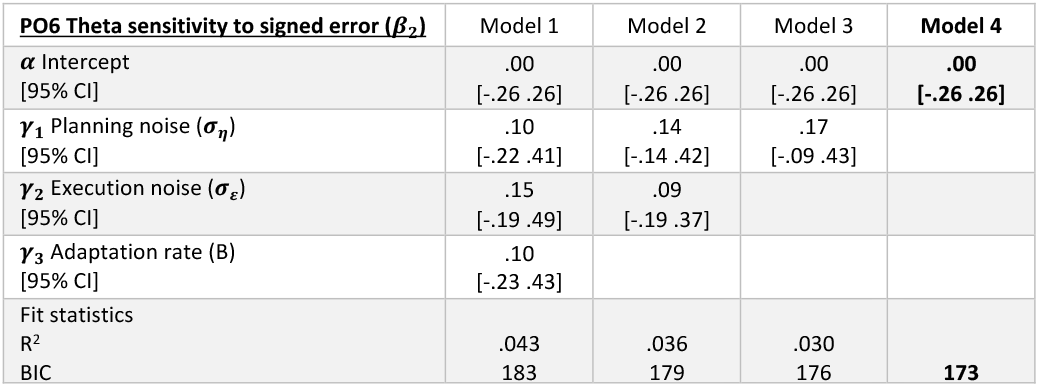
Multivariate linear model on the participant level (6U participants), with right occipital (PO6) theta sensitivity to the signed error as the independent variable (Supplementary Table 1, Model PO6). In order to compare the strength of the correlations, all variables were z-score normalized. Stepwise removal of variables with the smallest effect size was continued as long as the Bayesian Information Criterion decreased, and the last model for which this criterion held was selected. Individual differences in theta sensitivity to the signed error were not explained by planning noise, execution noise and adaptation rate.

| **PO6 Theta sensitivity to signed error ($\beta_2$)** | Model 1 | Model 2 | Model 3 | **Model 4** |
| --- | --- | --- | --- | --- |
| $\alpha$ Intercept<br>[95% CI] | .00<br>[-.26 .26] | .00<br>[-.26 .26] | .00<br>[-.26 .26] | **.00**<br>**[-.26 .26]** |
| $\gamma_1$ Planning noise ($\sigma_\eta$)<br>[95% CI] | .10<br>[-.22 .41] | .14<br>[-.14 .42] | .17<br>[-.09 .43] | |
| $\gamma_2$ Execution noise ($\sigma_\varepsilon$)<br>[95% CI] | .15<br>[-.19 .49] | .09<br>[-.19 .37] | | |
| $\gamma_3$ Adaptation rate (B)<br>[95% CI] | .10<br>[-.23 .43] | | | |
| Fit statistics |  |  |  |  |
| R<sup>2</sup> | .043 | .036 | .030 |  |
| BIC | 183 | 179 | 176 | **173** |

## SUPPLEMENTARY FIGURE LEGENDS

**Supplementary Figure 1:**
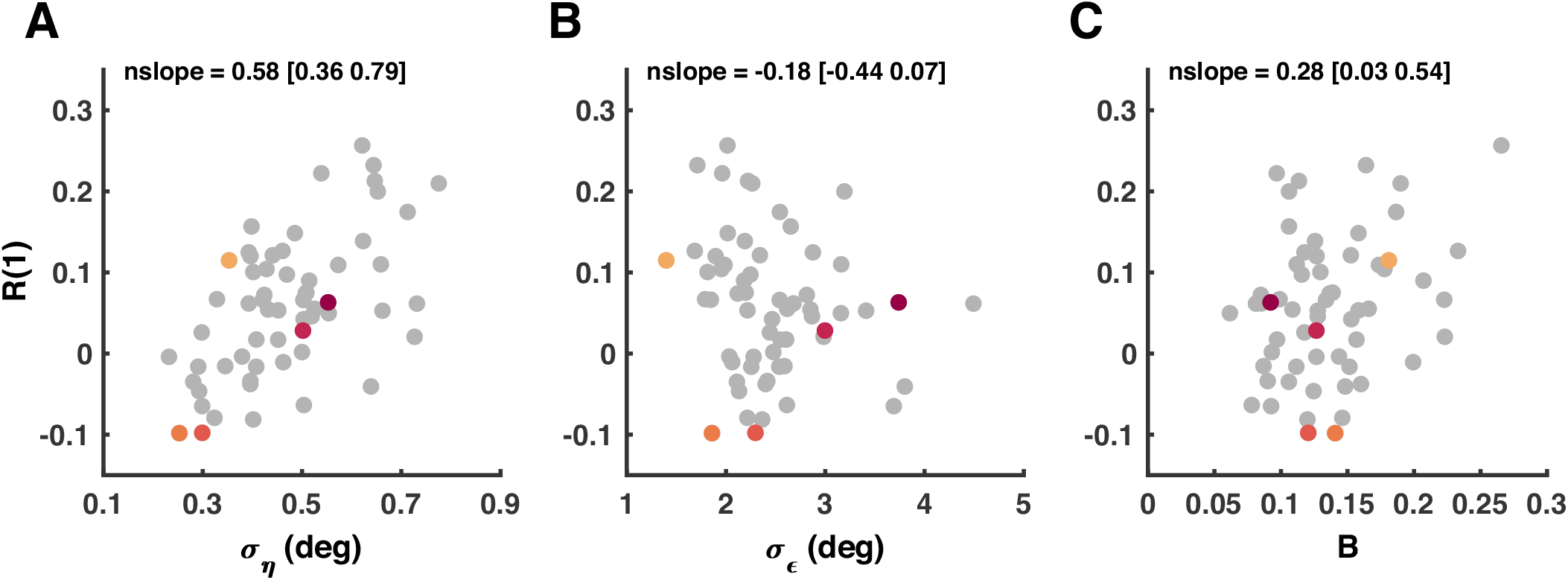
Relations between planning (09), execution noise (*σ_ε_*), adaptation rate (B) and the lag-1 autocorrelation in the baseline block R(1). Dots represent the absolute values and ‘nslope’ represents the linear relation of the z-score normalized values (slope [95%CI]).

**Supplementary Figure 2:**
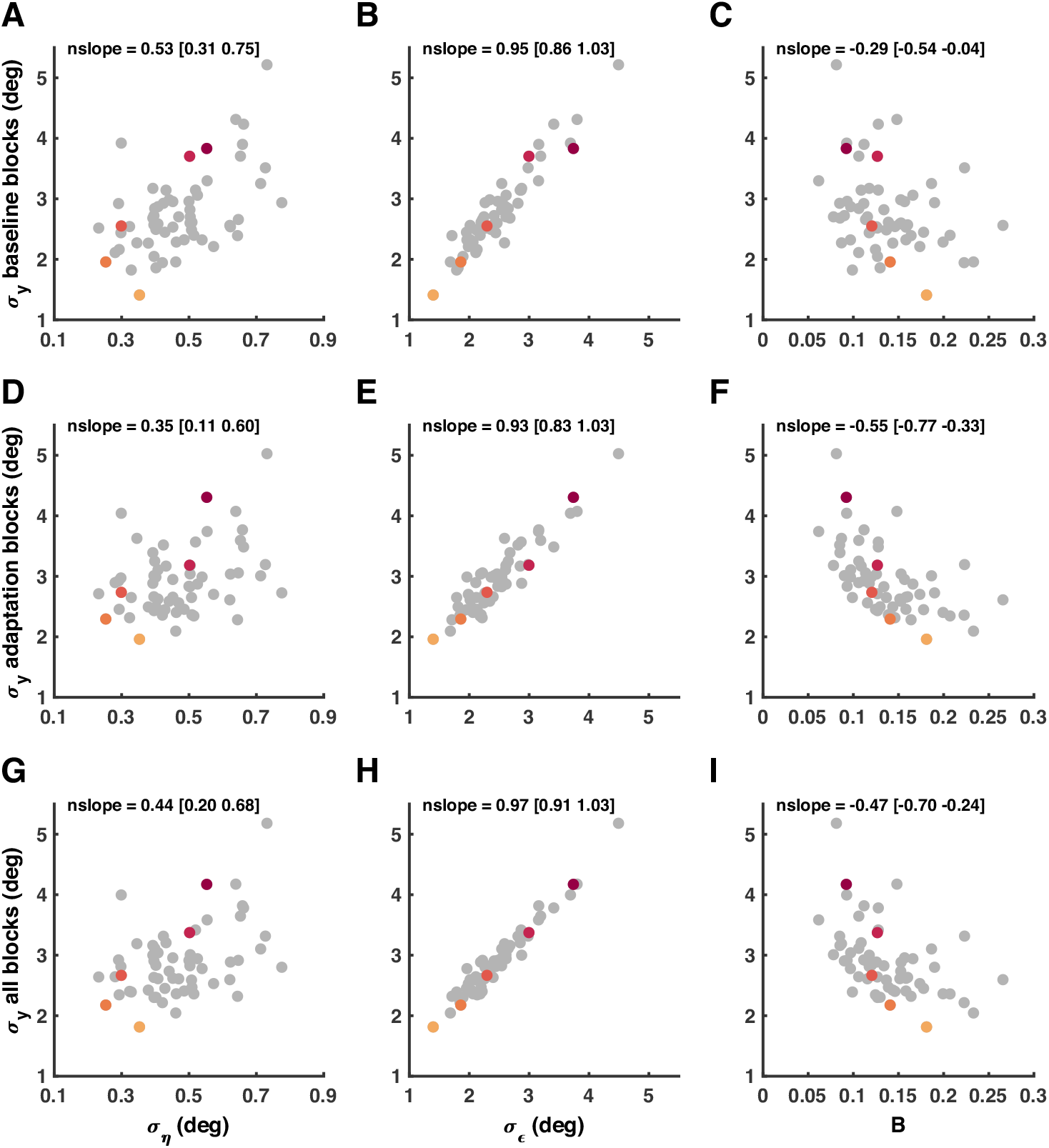
Relations between planning (09), execution noise (*σ_ε_*), adaptation rate (B) and the standard deviation of signed errors o=[s] in the baseline block **(A-C)**, the adaptation block **(D-F)** and both blocks combined **(G-I)**. Dots represent the absolute values and ‘nslope’ represents the linear relation of the z-score normalized values (slope [95%CI]).

**Supplementary Figure 3:**
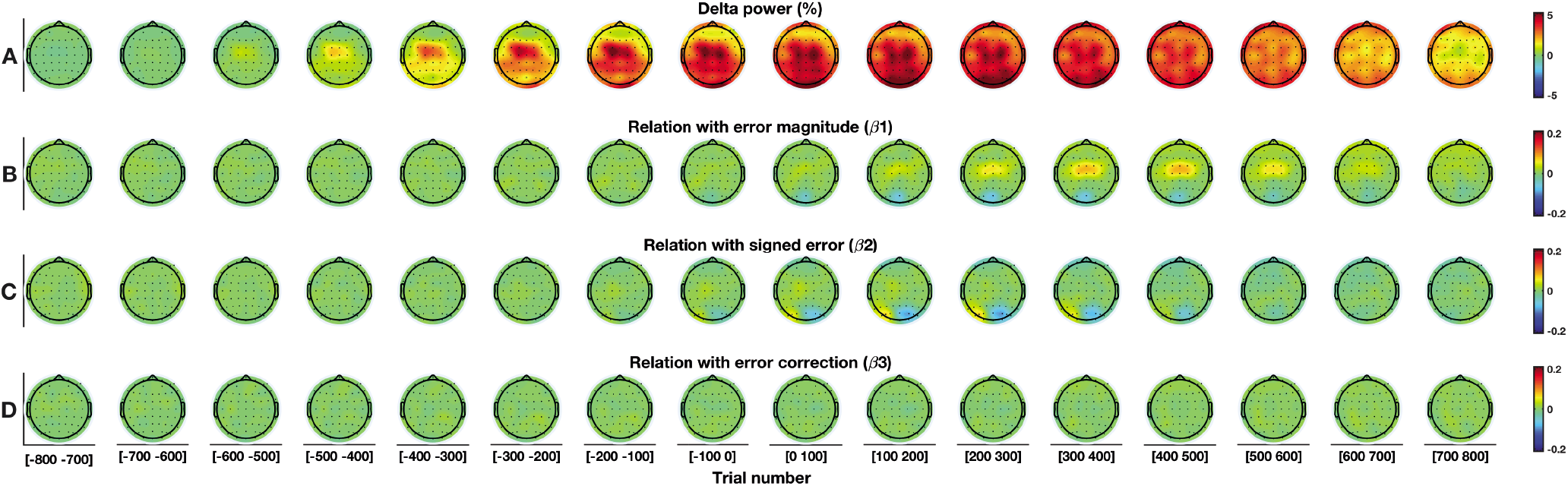
Trial level analysis in the delta frequency band (2-3Hz). For this figure, the full mixed-effects model is used (Eq. 6) **A:** Delta power. **B:** Relation with error magnitude. **C:** Relation with signed error. **D:** Relation with error correction.

**Supplementary Figure 4:**
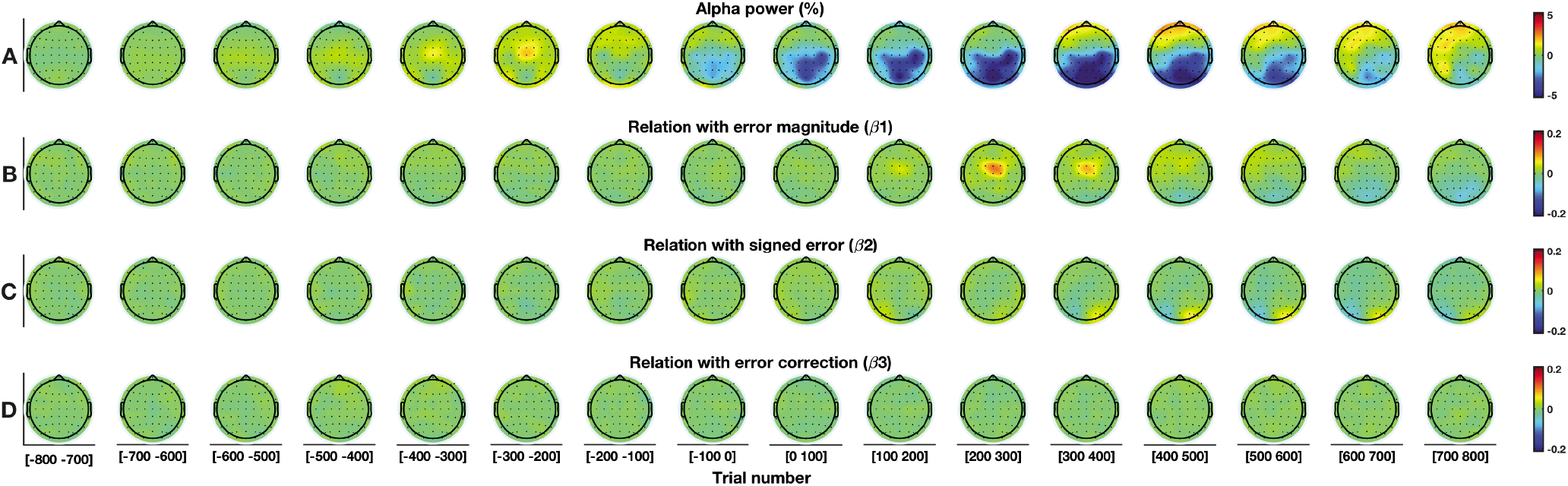
Trial level analysis in the alpha frequency band (9-14Hz). For this figure, the full mixed-effects model is used (Eq. 6) **A:** Alpha power. **B:** Relation with error magnitude. **C:** Relation with signed error. **D:** Relation with error correction.

**Supplementary Figure 5:**
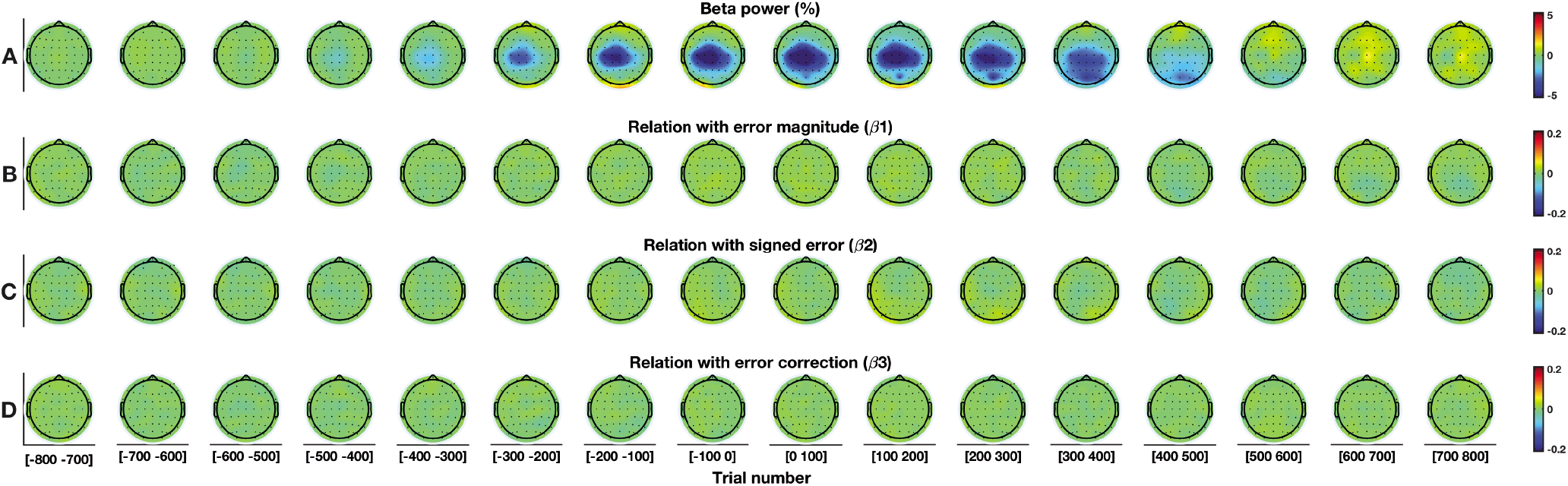
Trial level analysis in the beta frequency band (15-30Hz). For this figure, the full mixed-effects model is used (Eq. 6) **A:** EEG amplitude. **B:** Relation with error magnitude. **C:** Relation with signed error. **D:** Relation with error correction.

**Supplementary Figure 6:**
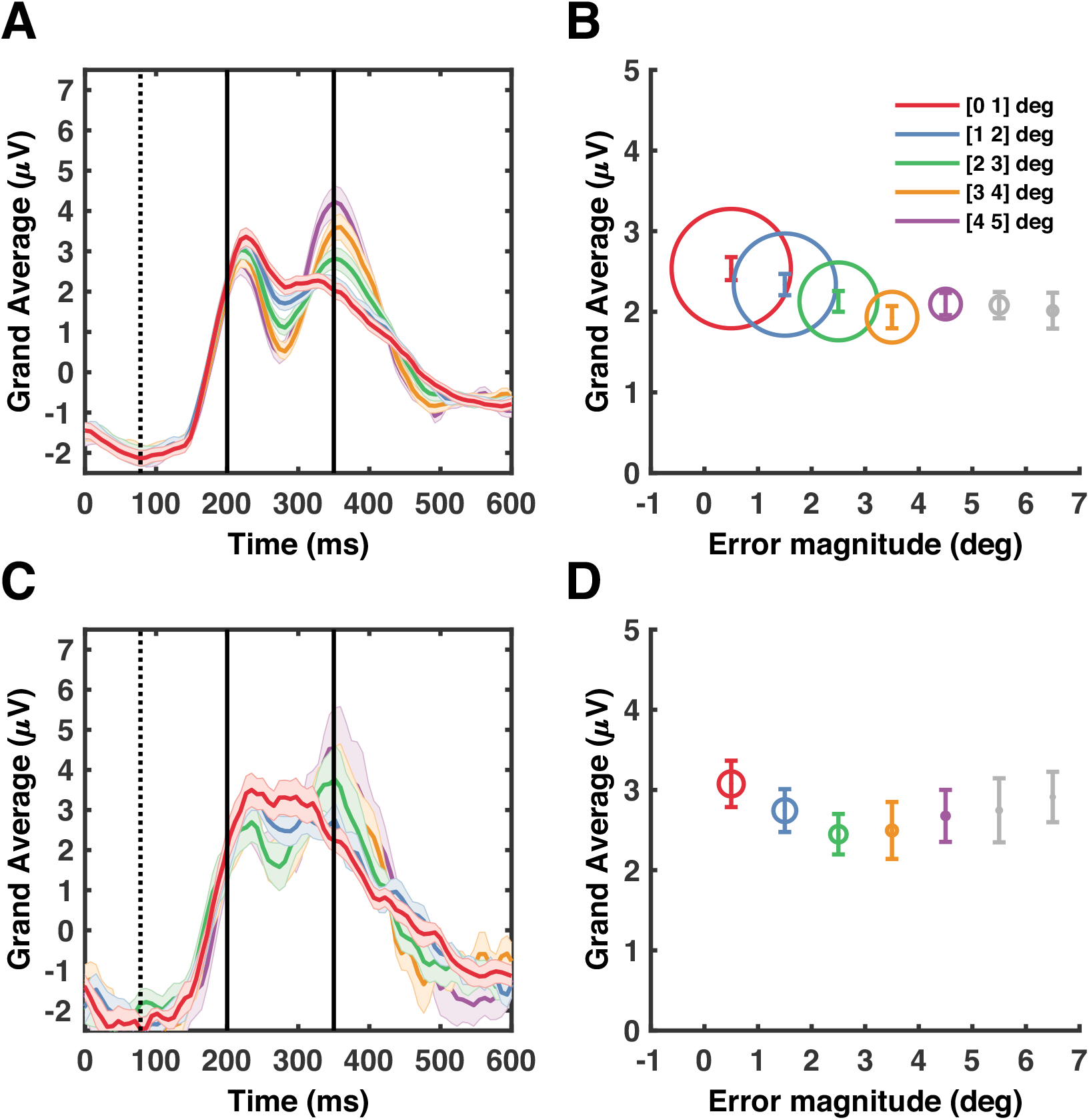
Error sensitivity in channel FCz in the time domain. For visualization purposes, trials have been binned and averaged based on error magnitude. Black dotted lines represent the movement end (78ms), relative to the onset of visual feedback. Black solid lines represent a post-feedback window of 200-350ms (Palidis et al., 2019). **A:** Average EEG amplitude across subjects. Error bars represent the standard error of the mean. The average EEG amplitude shows a wave-like signal within a larger positive response. Note that the post-feedback window contains both the negative and a part of the positive peak of the wave-like signal. **B:** Average EEG amplitude in the in the post-feedback window. The diameter of the circles is related to the average number of trials in the bin. **C:** Similar to panel A, but with a smaller sample. For visualization purposes, only the first 20 participants were included and of these participants only the first 150 trials of the adaptation block were included. With a smaller sample size the wave-like signal within the large positive response becomes less visible. **D:** Similar to panel B, but with the smaller sample. Note that the average EEG amplitude in the post-feedback window (200-350ms) appears similar for small and large error magnitudes.

**Supplementary Figure 7:**
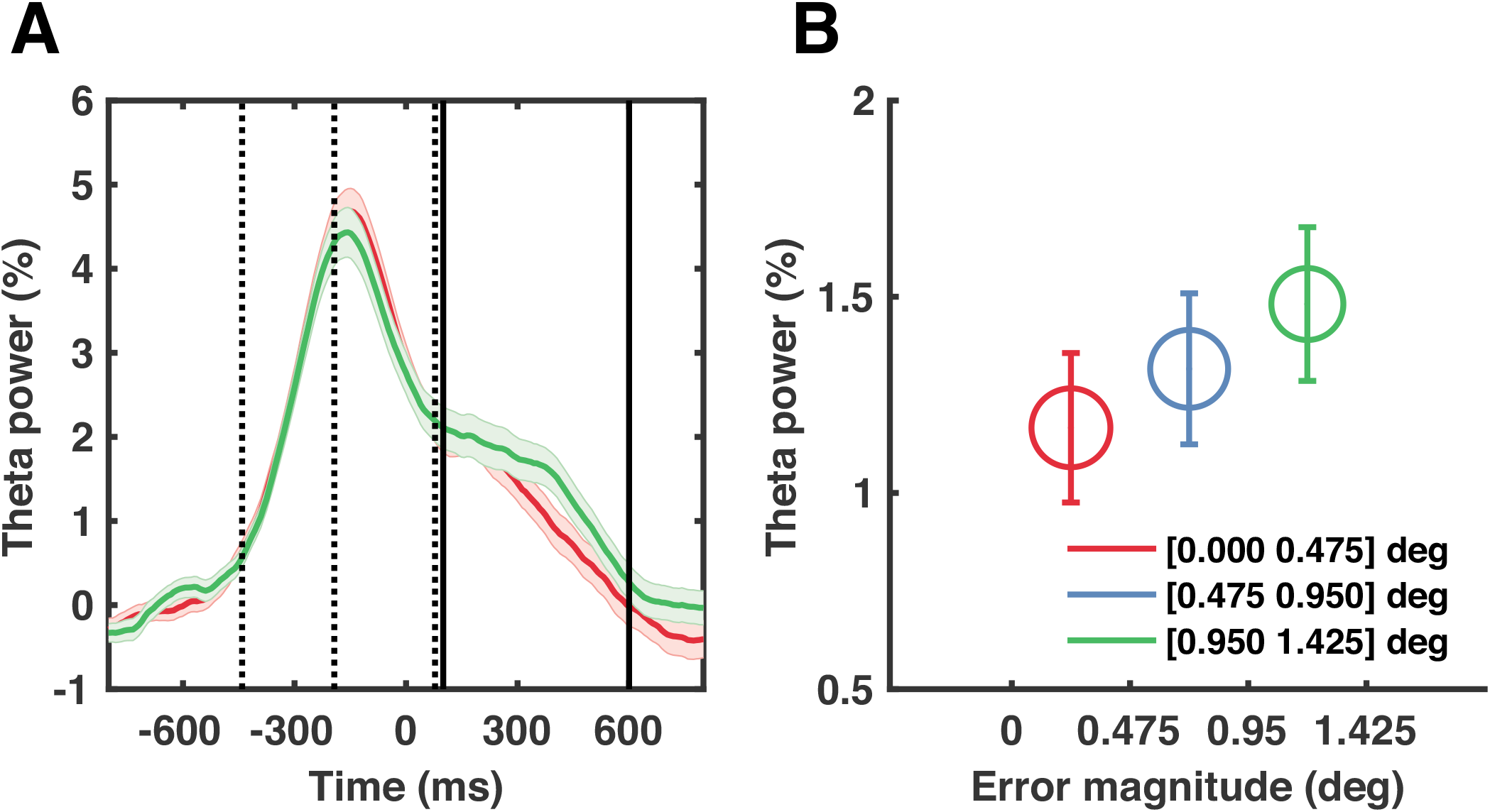
Error sensitivity in channel FCz in the subset of trials that were completely inside the target (error magnitude < 1.43). For visualization purposes, trials have been binned and averaged based on error magnitude. **A:** Average theta power (4-8Hz) across subjects. Error bars represent the standard error of the mean. Black dotted lines represent the average target onset (−441ms), movement onset (−193ms) and movement end (78ms), relative to the onset of visual feedback. Black solid lines represent the post-feedback window (100-600ms). **B:** Average theta power in the in the post-feedback window. The diameter of the circles is related to the average number of trials in the bin.

**Supplementary Figure 8:**
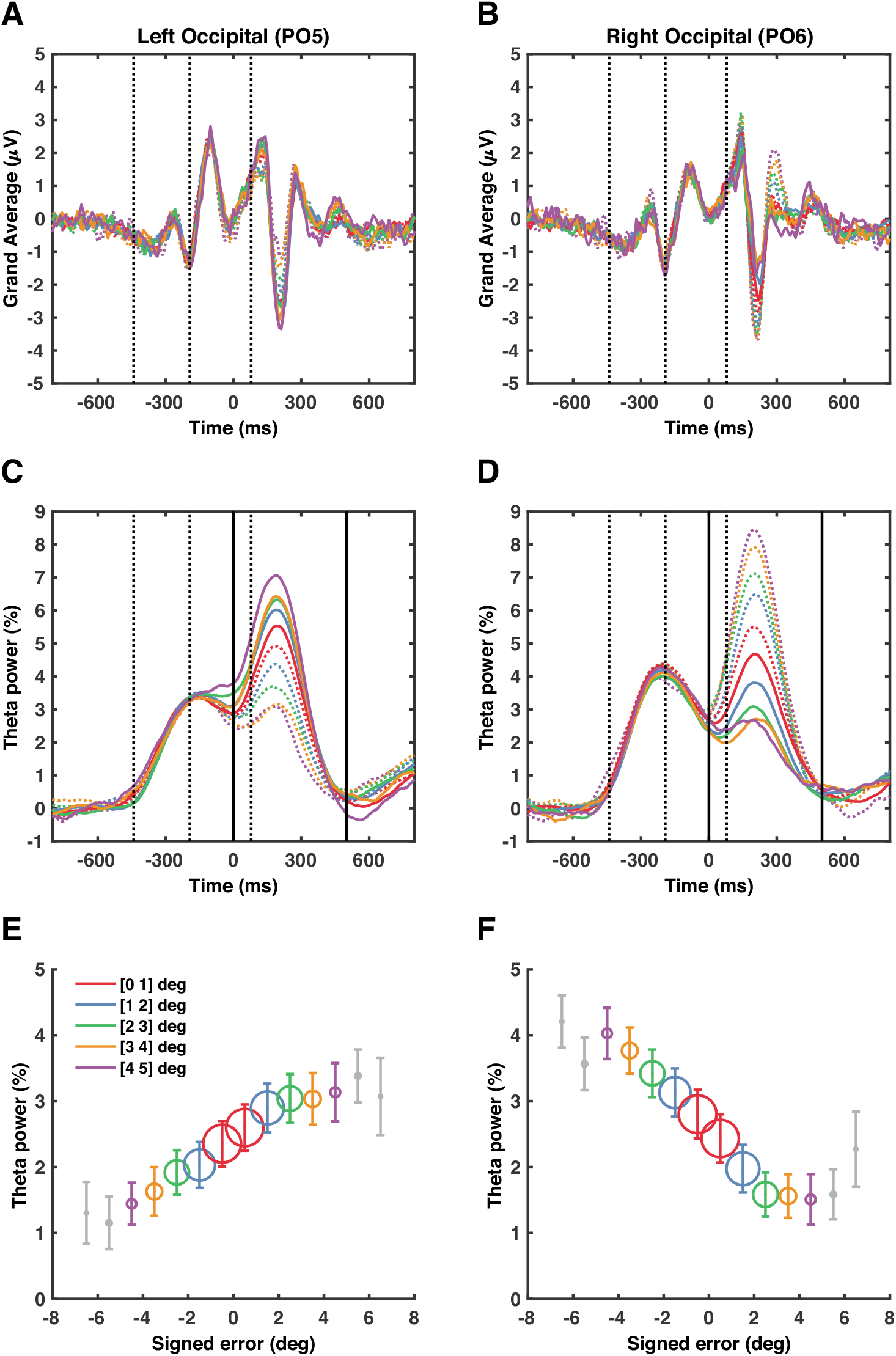
Sensitivity to the signed error in channel PO5 (left column) and PO6 (right column). For visualization purposes, trials have been binned and averaged based on signed error. Colored dotted lines represent bins with a negative sign. Black dotted lines represent the average target onset (−441ms), movement onset (−193ms) and movement end (78ms), relative to the onset of visual feedback. Black solid lines represent the post-feedback window (0-500ms). **A,B:** Average EEG amplitude across subjects. **C,D:** Average theta power (4-8Hz) across subjects. **E,F:** Average theta power in the post-feedback window. The diameter of the circles is related to the average number of trials in the bin.

**Supplementary Figure 9:**
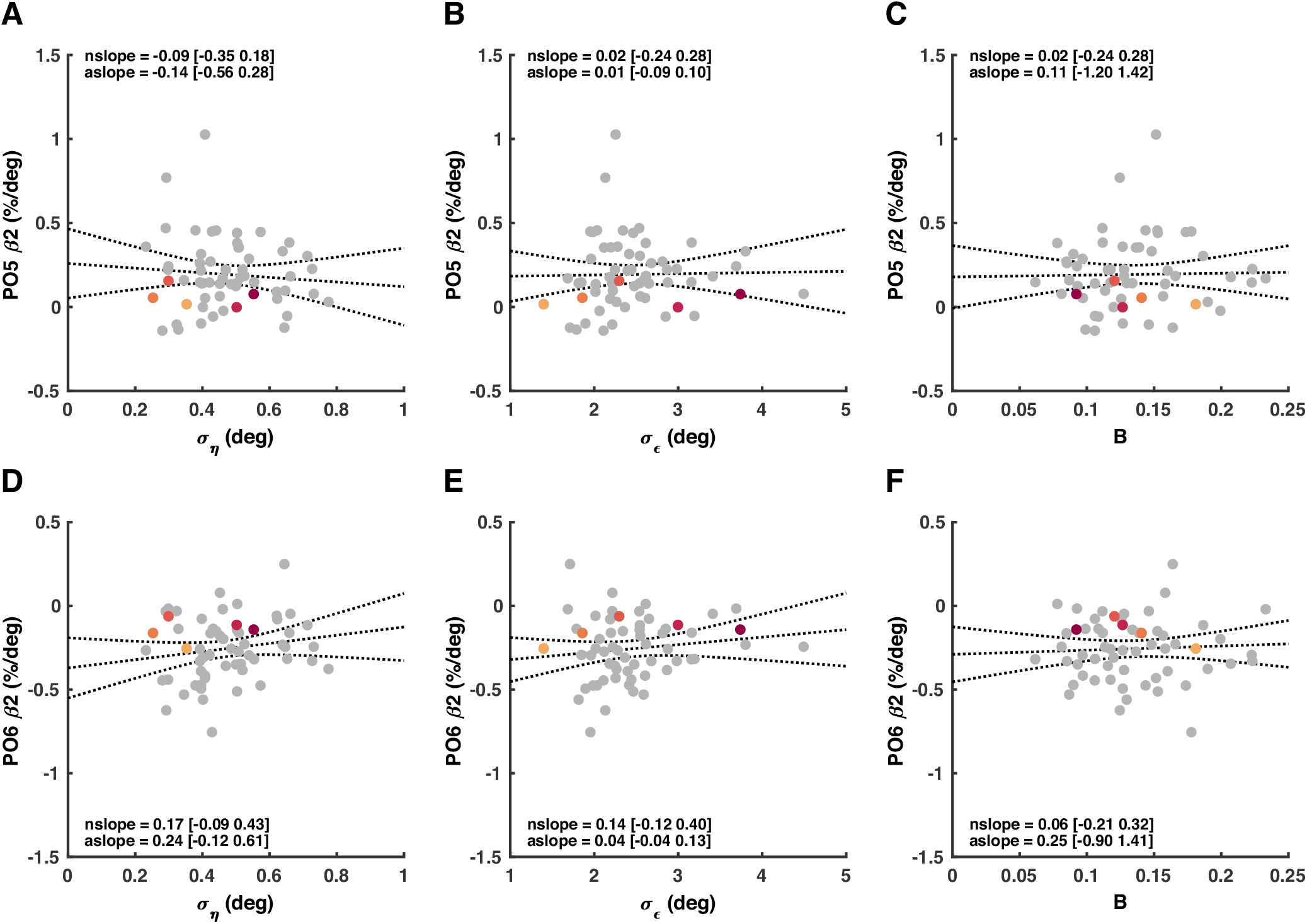
Linear model on the participant level (60 participants), with occipital theta (4-8Hz) sensitivity to the signed error as the dependent variable (*β*_2_) (Supplementary Table 1). **A-C:** Left occipital (PO5) theta sensitivity to the signed error was not related to planning (09 [s]), execution noise (*σ_η_*[*s*]), nor adaptation rate (B) (Supplementary Table 2). **D-F:** Right occipital (PO6) theta sensitivity to the signed error was also not related to planning (09 [s]), execution noise (*σ_ε_*[*s*]), nor adaptation rate (B) (Supplementary Table 3). Black lines represent the absolute slope (‘aslope’) and the confidence interval. The z-score normalized slope is also shown (‘nslope’).

